# Restoring the Chondron: Pericellular Matrix Reconstitution Enhances Mechano-Inflammatory Resilience and Modulates Chondrocyte–Neuron Crosstalk

**DOI:** 10.64898/2026.09.09.750375

**Authors:** Huan Meng, Junxuan Ma, Jiangyao Xu, Olivier Chassande, Line Kawtharany, Martin J. Stoddart, Sibylle Grad, Zhen Li

## Abstract

Chondrocytes in native articular cartilage are enclosed within a collagen VI (COL VI)-rich pericellular matrix (PCM), forming functional units known as chondrons. However, enzymatic isolation disrupts the PCM, and the functional consequences of its loss and restoration remain poorly understood.

In this study, primary human chondrocytes were cultured in alginate beads to promote PCM reconstitution and subsequently recovered as reconstituted chondrons. Chondrocytes and reconstituted chondrons were encapsulated in gelatin methacryloyl hydrogels and compared under inflammatory and mechanical stimulation. Alginate preconditioning generated chondron-like units with a distinct COL VI-positive PCM that was retained after transfer to three-dimensional culture. Under inflammatory conditions, reconstituted chondrons exhibited reduced inflammatory, catabolic, neuroinflammatory, and angiogenic responses compared with isolated chondrocytes at both gene and protein levels. Under interleukin-1β stimulation, mechanical loading further increased inflammatory gene expression in chondrocytes in a donor-dependent manner, whereas responses remained comparatively limited in reconstituted chondrons. Conditioned medium from reconstituted chondrons was also associated with lower capsaicin-and potassium chloride-evoked calcium responses in human induced pluripotent stem cell-derived sensory neurons than corresponding chondrocyte-conditioned medium.

These findings demonstrate that reconstitution of a COL VI-rich PCM restores a chondron-like pericellular microenvironment that attenuates inflammatory activation and buffers load-associated inflammatory amplification, while exploratory sensory neuron experiments suggest downstream modulation of neuronal responsiveness. PCM restoration may therefore provide a promising strategy for cartilage tissue engineering and regenerative applications.

**Graphical abstract:** 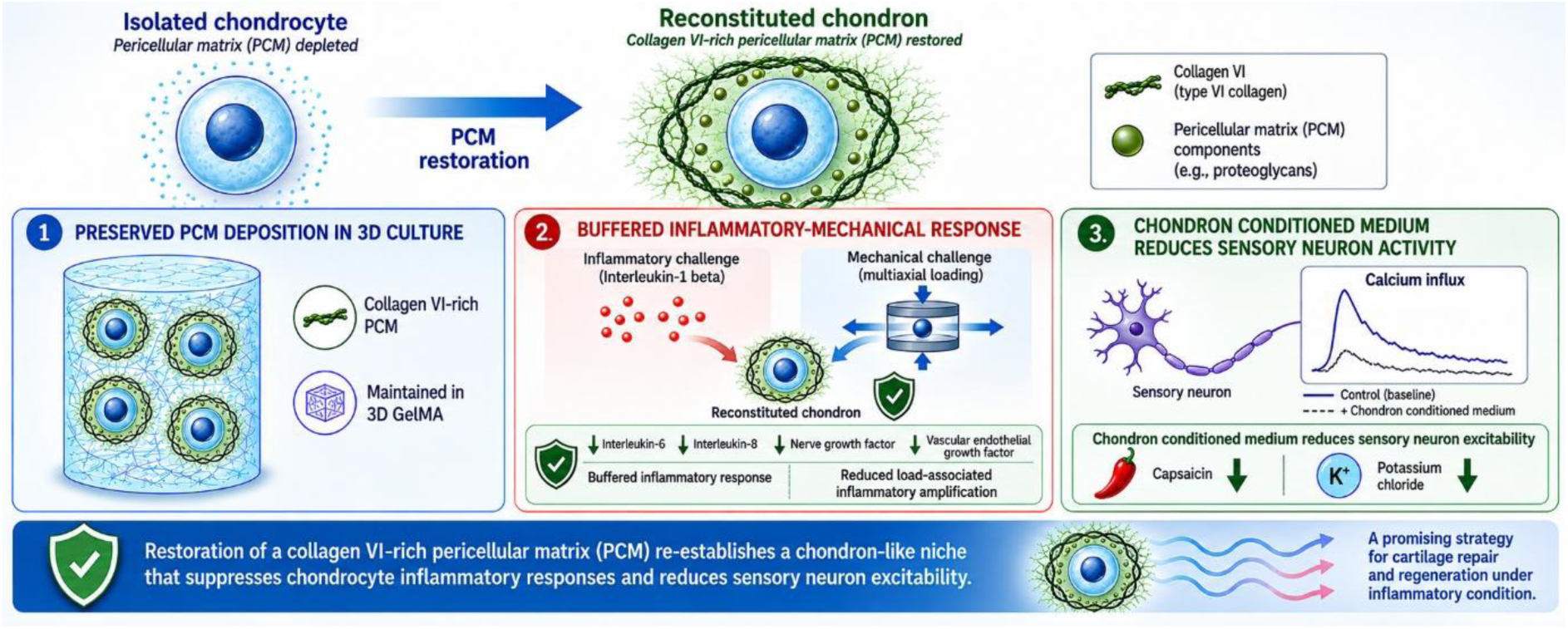

## 1. Introduction

Articular cartilage is a highly specialized avascular connective tissue that provides a low-friction, load-bearing surface in synovial joints[1]. Chondrocytes represent the predominant resident cell population and are principally responsible for maintaining extracellular matrix homeostasis[2]. However, chondrocytes do not exist in isolation. Instead, they are enclosed within a distinct pericellular matrix (PCM), forming a structural and functional unit termed the chondron[3, 4]. Compositionally distinct from the surrounding extracellular matrix, the pericellular matrix (PCM) is enriched in collagen type VI (COL VI) and associated matrix molecules and serves as the immediate microenvironment of the cell[5]. Increasing evidence suggests that the PCM regulates cellular behavior through its roles in mechanical filtering[6], mechanotransduction[7, 8], molecular transport[6], growth factors release[9] and matrix homeostasis[5]. Together, these observations suggest that the chondron, rather than the isolated chondrocyte, may represent the biologically relevant unit through which cartilage senses and responds to its environment.

Articular cartilage exhibits a limited intrinsic capacity for repair, motivating the development of a range of cell-based cartilage regeneration strategies[10]. Autologous chondrocyte implantation (ACI)[11], matrix-assisted ACI (MACI)[12], and numerous tissue-engineering approaches[13] rely on primary chondrocytes enzymatically isolated and expanded in vitro. However, enzymatic isolation disrupts the native PCM, resulting in the loss of the chondron architecture prior to implantation[14]. Subsequent monolayer expansion further separates chondrocytes from their native pericellular environment and may contribute to phenotypic alterations during culture[15]. Consequently, current cartilage repair strategies primarily seek to restore the cellular component of cartilage while overlooking the protective microenvironment that normally accompanies it. Whether this loss of the PCM represents a critical limitation of current cartilage-repair strategies remains unknown.

As the immediate interface between the cell and the surrounding extracellular matrix, the PCM can positioned to regulate how biochemical and mechanical signals are transmitted to the cell[16–18]. Alterations in this pericellular environment, including those observed in degenerative cartilage such as osteoarthritis[19], may therefore modify how chondrocytes respond to inflammatory and mechanical stress[20] and contribute to disturbed matrix homeostasis[17]. Importantly, inflammatory and mechanical stimuli act concurrently within the joint and interact bidirectionally[21, 22], yet whether PCM integrity regulates this mechano-inflammatory crosstalk remains poorly understood. Changes in chondrocyte signaling may also reshape the cartilage secretome, including mediators implicated in sensory-neuron sensitization and pain[23]. Emerging evidence further suggests that cartilage-derived factors can influence sensory-neuron responsiveness[24], indicating that alterations in the chondrocyte microenvironment may affect not only local tissue homeostasis but also intercellular communication within the joint. Collectively, these observations suggest that PCM is an important regulator of how chondrocytes integrate mechanical and biochemical cues.

Despite the recognized importance of PCM in native cartilage, it remains unclear whether the protective functions of the chondron can be reconstructed following enzymatic isolation and expansion of primary chondrocytes. Specifically, it is unknown whether a regenerated PCM can be maintained within a 3D tissue-engineering environment, whether it modifies cellular responses to combined inflammatory and mechanical stimulation, and whether PCM-dependent changes in the cartilage secretome influence sensory-neuron responsiveness.

Therefore, the aim of this study was to reconstruct a chondron-like cellular unit by promoting PCM formation around isolated primary human chondrocytes and to determine whether PCM restoration improves cellular resilience within an inflammatory and mechanically active 3D environment. We hypothesized that reconstituted chondrons would maintain COL VI-rich PCM and exhibit attenuated inflammatory responses and reduced loading-associated amplification of inflammatory signaling compared with isolated chondrocytes. As an exploratory extension, we further examined whether conditioned media generated under these conditions produced distinct effects on sensory-neuron calcium responsiveness.

## 2. Materials and methods

### 2.1 Cell isolation and culture

Three culture media were used throughout the study. Expansion medium consisted of high-glucose Dulbecco’s Modified Eagle Medium (DMEM-HG, REF 52100-021, Gibco, USA) supplemented with 10% fetal bovine serum (FBS, REF 10500-064, Gibco, USA), 1% non-essential amino acids (NEAA, REF 11140-035, Gibco, USA), 1% penicillin/streptomycin (P/S; REF 15140-122, Gibco, USA), 1 ng/mL human recombinant transforming growth factor (TGF-β1, REF 30R-AT072, Fitzgerald, USA), and 5 ng/mL basic fibroblast growth factor (bFGF, REF 30R-AF015, Fitzgerald, USA). Chondrogenic medium contained high-glucose DMEM, 1% NEAA, 1% P/S, 1% ITS Premix (REF 354352, Corning, USA), 50 μg/mL L-ascorbic acid 2-phosphate (REF A8960-5G, Sigma-Aldrich, USA), 0.1 μM dexamethasone (REF D2915, Sigma-Aldrich, USA), and 10 ng/mL TGF-β1. Chondropermissive medium consisted of high-glucose DMEM supplemented with 1% ITS Premix, 50 μg/mL L-ascorbic acid 2-phosphate, 1% NEAA, and 1% P/S.

Human articular chondrocytes were isolated from osteoarthritic knee cartilage obtained during total knee arthroplasty with informed consent at Spital Davos (see Supplementary Table S1 for donor details). The Swiss Human Research Act does not apply to research which involves anonymized biological material. Therefore, this project does not need to be approved by the ethics committee. Chondrocytes were obtained from 3 independent human donors. The clinical characteristics of human donors are summarized in Supplementary Table S1. Cartilage was minced and digested overnight at 37°C with collagenase type II[25] (2 mg/mL, 450 U/mL; REF LS004177, Worthington Biochemical Corporation, USA). Isolated cells were filtered through a 40 μm cell strainer (REF 431750, Corning, USA), expanded in T300 flasks (1 × 10⁶ cells/flask) using expansion medium, and passaged once. Passage 1 (P1) chondrocytes were used for all subsequent experiments.

Human native chondrons were isolated from osteoarthritic knee cartilage using an established enzymatic digestion protocol with minor modifications[26]. Briefly, cartilage fragments were digested overnight in DMEM containing 225 U/mL collagenase type II (REF LS004177, Worthington Biochemical Corporation, USA), and isolated chondrons were recovered by filtration through a 70 μm cell strainer. Freshly isolated native chondrons were fixed immediately for immunofluorescence characterization.

### 2.2 PCM reconstitution and 3D GelMA culture

Chondrocytes were seeded in sterile alginate beads at a density of 5 × 10⁶ cells/mL. Alginate was reconstituted at 1.2% w/v (Alginic acid sodium, Cat. No A1112 Sigma-Aldrich, USA) and beads were formed by extrusion through a 21-gauge needle into 102 mM CaCl₂ solution and crosslinked for approximately 10 min. Cell-laden alginate beads were cultured in chondrogenic medium for 14 days to promote the reconstitution of a collagen type VI-rich pericellular matrix. Following culture, alginate beads were dissolved using 55 mM sodium citrate (S4641, Sigma, USA), and the recovered cells, hereafter referred to as reconstituted chondrons, were collected for subsequent experiments.

For three-dimensional culture, chondrocytes or reconstituted chondrons were suspended in 5% (w/v) gelatin methacryloyl (GelMA; gel strength 300 Bloom, degree of substitution 60%; Cat. No. 900622-1G, Advanced BioMatrix, USA) containing 0.1% (w/v) lithium phenyl-2,4,6-trimethylbenzoylphosphinate (LAP, 900889, Sigma-Aldrich, USA) photoinitiator at a density of 5 × 10⁶ cells/mL. Cell-laden hydrogel cylinders with a diameter of 8 mm and height of 4 mm, were casted in customized moulds photocrosslinked by blue-light irradiation (LunaX™ Crosslinker, Gelomics, Australia) for 5 min and cultured in chondropermissive medium for 24 h before downstream experiments.

### 2.3 Multiaxial mechanical loading and inflammatory stimulation

A custom-designed MultiWell bioreactor developed at the AO Research Institute Davos was used to apply controlled multiaxial mechanical stimulation to three-dimensional GelMA constructs[27]. The bioreactor accommodates 16 samples simultaneously within a sterile culture chamber and applies compressive and rotational movements through ceramic loading contacts positioned above each construct. Individual loading units are independently motor-controlled and integrated with load cells for continuous force monitoring throughout the loading period. Samples were maintained under standard cell culture conditions (37°C, 5% CO₂) during loading. A schematic overview of the bioreactor and loading configuration is provided in Figure 5A.

Mechanical loading was performed in chondropermissive medium using a physiologically relevant multiaxial loading protocol designed to mimic the combined compression and shear experienced by articular cartilage during joint motion. GelMA constructs were subjected to loading at 1 Hz for 1 h per day, consisting of 10–20% dynamic compressive strain superimposed on 10% static offset strain and 0–12° oscillatory shear loading[28]. Unloaded control samples were cultured under identical conditions in chondropermissive medium in the sample holders, without mechanical stimulation. The unloaded control samples and the loaded samples were all treated +/-1 ng/mL IL-1β (200-01B-10UG, Thermo Fisher, USA). Culture media and/or IL-1β was refreshed every day for 7 days,

### 2.4 Histology, immunofluorescence and microscopy

For histological evaluation, GelMA constructs were fixed in 4% neutral-buffered formalin for 24 h, embedded in paraffin, and sectioned at 5 μm thickness. Sections were stained with 0.1% Safranin O/ 1% Fast Green using standard protocols to evaluate glycosaminoglycan deposition. Brightfield images were acquired using an Olympus BX63 microscope (Olympus Corporation, Japan).

For immunofluorescence staining, GelMA sections were immunolabelled for collagen type VI (COL VI) to visualize the pericellular matrix, with nuclei counterstained using DAPI. iPSC-derived sensory neurons were fixed with 4% paraformaldehyde for 15 min at room temperature and immunostained for βIII-tubulin (TUBB3) and calcitonin gene-related peptide (CGRP) to assess neuronal differentiation and sensory neuronal phenotype. Primary and secondary antibodies are listed in Supplementary Table S2.

Fluorescence images were acquired using a Zeiss LSM700 laser scanning confocal microscope (Carl Zeiss, Germany). For sensory neuron imaging, z-stack images were acquired at 5 μm intervals over three sections, followed by maximum-intensity projection for image analysis. Identical acquisition settings were maintained for all samples within each experiment.

### 2.5 Gene expression analysis

Total RNA was isolated from chondrocytes and chondrons cultured in GelMA hydrogels. Prior to RNA extraction, GelMA constructs were enzymatically digested with 450 U/mL collagenase type II (REF LS004177, Worthington Biochemical Corporation, USA) for approximately 30 min at 37°C to facilitate hydrogel dissolution and improve RNA recovery. Cell suspensions were subsequently collected by centrifugation, and total RNA was extracted using the QIAwave RNA Mini Kit (Cat. No. 74534, QIAGEN, Germany) according to the manufacturer’s instructions.

RNA concentration and purity were determined using a NanoDrop One spectrophotometer (Thermo Fisher Scientific, USA). Equal amounts of RNA (500 ng) were reverse transcribed into complementary DNA (cDNA) using the SuperScript™ IV VILO™ Master Mix (Thermo Fisher Scientific, USA) following the manufacturer’s protocol.

Quantitative real-time PCR (RT-qPCR) was performed using TaqMan™ Gene Expression Assays (Thermo Fisher Scientific, USA) on a QuantStudio™ 7 Flex Real-Time PCR System (Applied Biosystems, Thermo Fisher Scientific, USA). Relative gene expression was calculated using the 2^^−ΔΔCt^ method, with RPLP0 used as the endogenous reference gene. All TaqMan assay IDs are listed in Supplementary Table S3.

### 2.6 Protein and DNA quantification

Culture supernatants were collected at the experimental endpoint, centrifuged to remove cellular debris, and stored at −80°C until analysis. The concentrations of human IL-6, IL-8, VEGF, and NGF were quantified using commercially available enzyme-linked immunosorbent assay (ELISA) kits according to the manufacturers’ instructions. Human IL-6 (DuoSet ELISA Development Kit, Cat. No. DY206-05, R&D Systems, USA), human IL-8 (DuoSet ELISA Development Kit, Cat. No. DY208-05, R&D Systems, USA), and human VEGF (DuoSet ELISA Development Kit, Cat. No. DY293B, R&D Systems, USA) were quantified using DuoSet ELISA kits, while human β-NGF was measured using a Human β-NGF SimpleStep ELISA Kit (Cat. No. ab193760, Abcam, UK). Samples and standards were analyzed in duplicate, and the mean absorbance was used for subsequent analyses. Absorbance was measured using a Tecan microplate reader (Tecan Group Ltd., Switzerland), and protein concentrations were calculated from standard curves generated according to the manufacturers’ instructions.

GelMA constructs were digested with 0.5 mg/mL proteinase K (cat. #1000144 Roche, Switzerland), and total DNA content was quantified using a Hoechst fluorescence assay (14530, Sigma, USA). Fluorescence was measured using the Tecan microplate reader.

### 2.7 iPSC-derived sensory neuron differentiation

Human induced pluripotent stem cells were obtained from Laboratoire BIOTIS INSERM U1026. The iPSCs were originally from the iPS-Quebec platform of the CHU de Québec-Université Laval research center. They were derived from 2 different human skin fibroblast cell lines and from one human peripheral blood mononuclear cell line. The iPSCs were reprogrammed with the CytoTuneTM-iPS 2.0 Sendai Repro-gramming Kit (Invitrogen, Carlsbad, California). iPSCs were differentiated into sensory neurons using an adapted protocol based on Maury et al.[29] and Muller et al.[30]. iPSCs were maintained on Geltrex-coated (REF A14133-02, Thermo Fisher Scientific, USA) culture plates, with Geltrex diluted 1:100 in DMEM/F12 according to the manufacturer’s recommendations. The morphology of differentiating iPSC-derived sensory neurons was monitored daily throughout the differentiation period using brightfield microscopy to assess neuronal outgrowth, network formation, and overall culture quality (Supplementary Figure S6).

For neural induction, iPSCs were cultured on Geltrex-coated 6-well plates and differentiated from day 0 to day 11 in a 1:1 mixture of DMEM/F12 and Neurobasal medium (REF 12348-017, Thermo Fisher Scientific, USA) supplemented with N2 (REF 17502-048, Thermo Fisher Scientific, USA), B27 (REF 12587-010, Thermo Fisher Scientific, USA), β-mercaptoethanol (REF 21985-023, Thermo Fisher Scientific, USA), L-ascorbic acid, Y-27632 (REF ab120129, Abcam, UK), alanyl-glutamine, MEM non-essential amino acids, trace elements A–C, SB431542 (REF S4317-5MG, Sigma-Aldrich, USA), LDN193189 (REF SML0559-5MG, Sigma-Aldrich, USA), CHIR99021 (REF SML1046-5MG, Sigma-Aldrich, USA), and gentamycin. SB431542 and LDN193189 were omitted after day 4, and CHIR99021 was removed from day 7 to initiate sensory neuron specification. DAPT (10 μM; REF S2215, Selleckchem, USA) was added from day 9 to day 12 to inhibit Notch signalling and promote neuronal differentiation (detailed media composition see Supplementary Table S4).

On day 11, immature sensory neurons were dissociated using Accutase (REF A11105-01, Thermo Fisher Scientific, USA), cryopreserved, and stored until use. For experiments, day-11 cells were thawed and seeded into 18-well μ-Slides (Cat. No. 81816-90, ibidi, Germany) to facilitate high-resolution immunofluorescence imaging and live-cell calcium imaging. Cells were further differentiated until day 18 in maintenance medium consisting of DMEM/F12 and Neurobasal (1:1) supplemented with 1× N2, 2% B27, 50 μg/mL L-ascorbic acid, 5 μM Y-27632, 20 ng/mL brain-derived neurotrophic factor (BDNF; REF 450-02, PeproTech, USA), 10 ng/mL glial cell line-derived neurotrophic factor (GDNF; REF 450-10, PeproTech, USA), 10 ng/mL nerve growth factor (NGF; REF 450-01, PeproTech, USA), 1X gentamycin, with medium changes every two days.

Differentiated sensory neurons (day 17) were treated with conditioned media for 24 h before calcium imaging and immunofluorescence analysis, which were performed on day 18.

### 2.8 Calcium imaging and analysis of neural response

Intracellular calcium influx was assessed in iPSC-derived sensory neurons following 24 h treatment with conditioned media using the fluorescent calcium indicator Fluo-4 AM (REF F14217, Thermo Fisher Scientific, USA). On day 18 of culture, neurons cultured in 18-well μ-Slides were washed once with Krebs-Ringer buffer (KB) and incubated for 30–60 min at 37°C with 5 μM Fluo-4 AM in KB containing 0.3% bovine serum albumin (BSA) and 1.25 mM probenecid (REF P36400, Thermo Fisher Scientific, USA). Following dye loading, cells were washed and incubated in fresh KB for 10 min before imaging.

Live-cell calcium imaging was performed in fresh KB at room temperature. During image acquisition, neurons were sequentially stimulated with 500 nM capsaicin (REF M2028, Sigma-Aldrich, USA) to activate transient receptor potential vanilloid 1 (TRPV1)-positive nociceptors, followed by 50 mM potassium chloride (KCl; REF P3911, Sigma-Aldrich, USA) to induce membrane depolarization and confirm neuronal viability (Figure 7B). Stimuli were added directly to the imaging chamber without interrupting image acquisition to minimize mechanical disturbance.

Fluorescence intensity was analyzed using Fiji software on the entire imaging field rather than individual neurons. Owing to the extensive neurite network and dense neuronal distribution following differentiation, reliable segmentation and tracking of individual neurons were not feasible. Therefore, changes in intracellular calcium were quantified as the normalized fluorescence intensity (ΔF/F₀) of the whole field of view, where F₀ represents the baseline fluorescence before stimulation and ΔF represents the change in fluorescence following stimulation. Peak ΔF/F₀ responses following capsaicin and KCl stimulation were used for subsequent statistical analyses.

### 2.9 Statistical analysis

Statistical analyses were performed using GraphPad Prism (GraphPad 11 Software, USA). Independent biological replicates represent human donors (N = 3), while experimental replicates are indicated as n = 9 where applicable. Data are presented as mean ± standard error of the mean (SEM) unless otherwise stated. Because chondrocytes and chondrons were from the same donors, paired statistical analyses were performed whenever appropriate. Comparisons involving multiple groups were analyzed using paired one-way or two-way repeated-measures analysis of variance (ANOVA) followed by Bonferroni’s multiple-comparison test, as appropriate for the experimental design. The exact statistical tests used for each experiment are indicated in the corresponding figure legends. Differences were considered statistically significant at P < 0.05.

## 3. Results

### 3.1 Reconstitution and characterization of the pericellular matrix in primary chondrocytes

To investigate the role of PCM in regulating inflammatory and mechanobiological responses, isolated chondrocytes and reconstituted chondrons were established as parallel experimental models, followed by culture in gelatin methacryloyl (GelMA) hydrogels under defined inflammatory and mechanical conditions (Fig. 1A). Native chondrons isolated directly from human articular cartilage exhibited the characteristic multicellular morphology surrounded by a collagen VI (COL VI)-positive pericellular matrix (Fig. 1B). Following alginate preconditioning, reconstituted chondrons displayed a comparable morphology as indicated by robust COL VI immunostaining (Fig. 1B). Quantitative analysis demonstrated that more than 90% of reconstituted chondrons were COL VI positive, comparable to native chondrons (Fig. 1C), confirming successful restoration of the pericellular matrix and validating the experimental model for subsequent mechanobiological studies.

**Figure 1.**
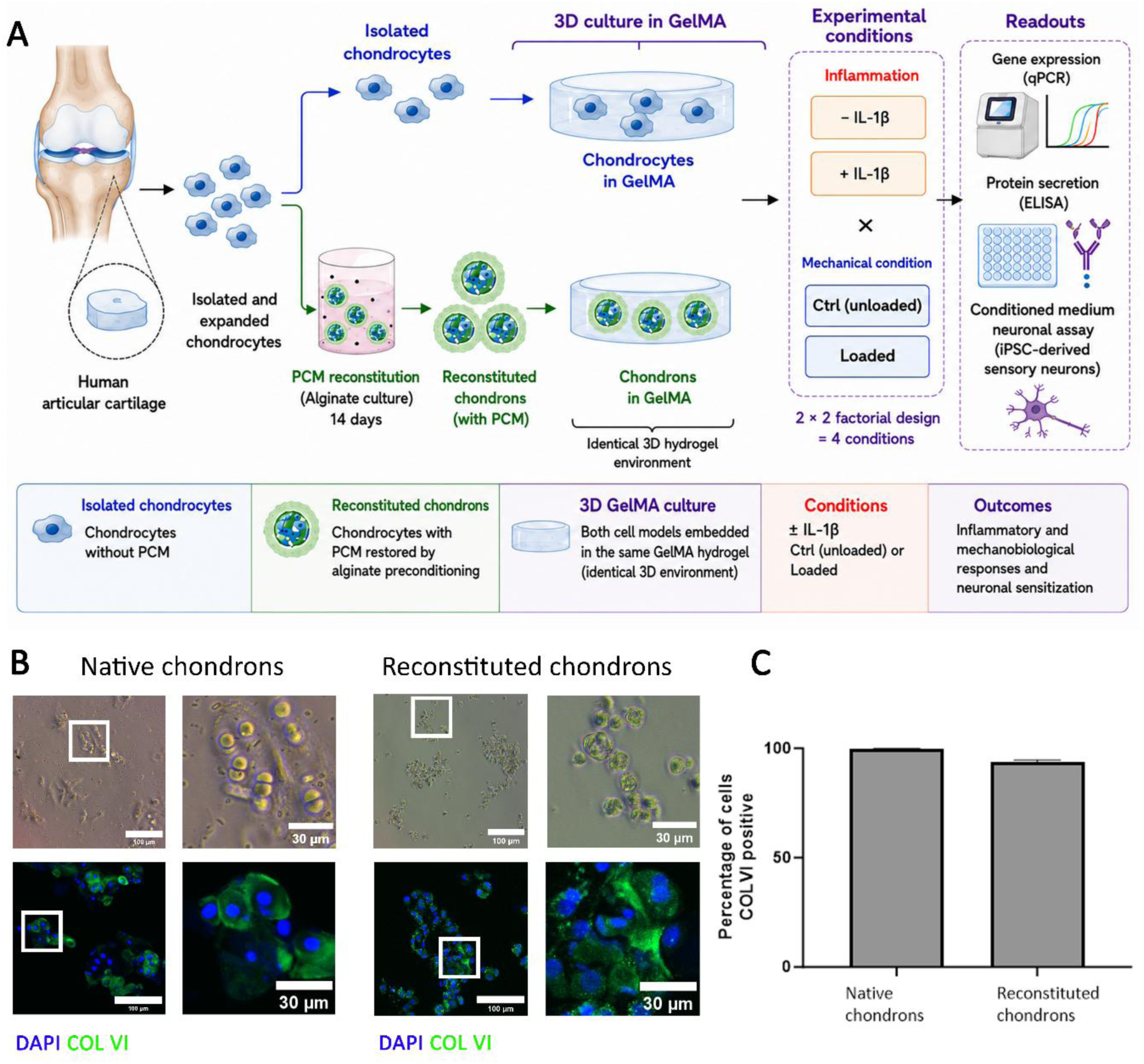
Reconstitution and characterization of the pericellular matrix in primary chondrocytes. (A) Schematic overview of the experimental workflow illustrating the generation of PCM-reconstituted chondrons from isolated primary human articular chondrocytes, followed by encapsulation in GelMA hydrogels and exposure to inflammatory (± IL-1β) and mechanical (Ctrl or Loaded) conditions for downstream analyses. (B) Representative bright-field and COL VI immunofluorescent staining images of native chondrons isolated directly from human articular cartilage and reconstituted chondrons following alginate preconditioning. (C) Quantification of percentage of COL VI-positive cells in native and reconstituted chondrons (n=6 images for each condition). Scale bars = 100 μm (low magnification) and 30 μm (high magnification). Bars represent mean ± SEM. N=3 donors while n=9 experimental replicates. Statistical analysis was performed using an unpaired Student’s t-test.

### 3.2 Reconstituted chondrons maintain pericellular matrix deposition and cartilage matrix synthesis in 3D GelMA culture

To determine whether the regenerated pericellular matrix was maintained following encapsulation in GelMA hydrogels, isolated chondrocytes and reconstituted chondrons were encapsulated in GelMA and cultured in a 3D environment for 7 days prior to further analyses (see section 2.2 and 2.3 for details). Immunofluorescence staining demonstrated minimal COL VI deposition around isolated chondrocytes, whereas reconstituted chondrons retained abundant COL VI surrounding the cells after 7 days of culture (Fig. 2A). Safranin-O/Fast green staining further demonstrated enhanced extracellular matrix deposition in reconstituted chondrons compared with chondrocytes (Fig. 2B). Consistent with these observations, expression of the cartilage matrix genes COL2 (P<0.05, Fig. 2D) and COMP (P<0.001, Fig. 2F) and PRG4 (P<0.05, Fig. 2E) was significantly higher in reconstituted chondrons, whereas ACAN expression remained comparable between the two groups (Fig. 2C). Together, these findings demonstrate that reconstituted chondrons maintain both the regenerated pericellular matrix and a matrix-producing chondrogenic phenotype during 3D GelMA culture.

**Figure 2.**
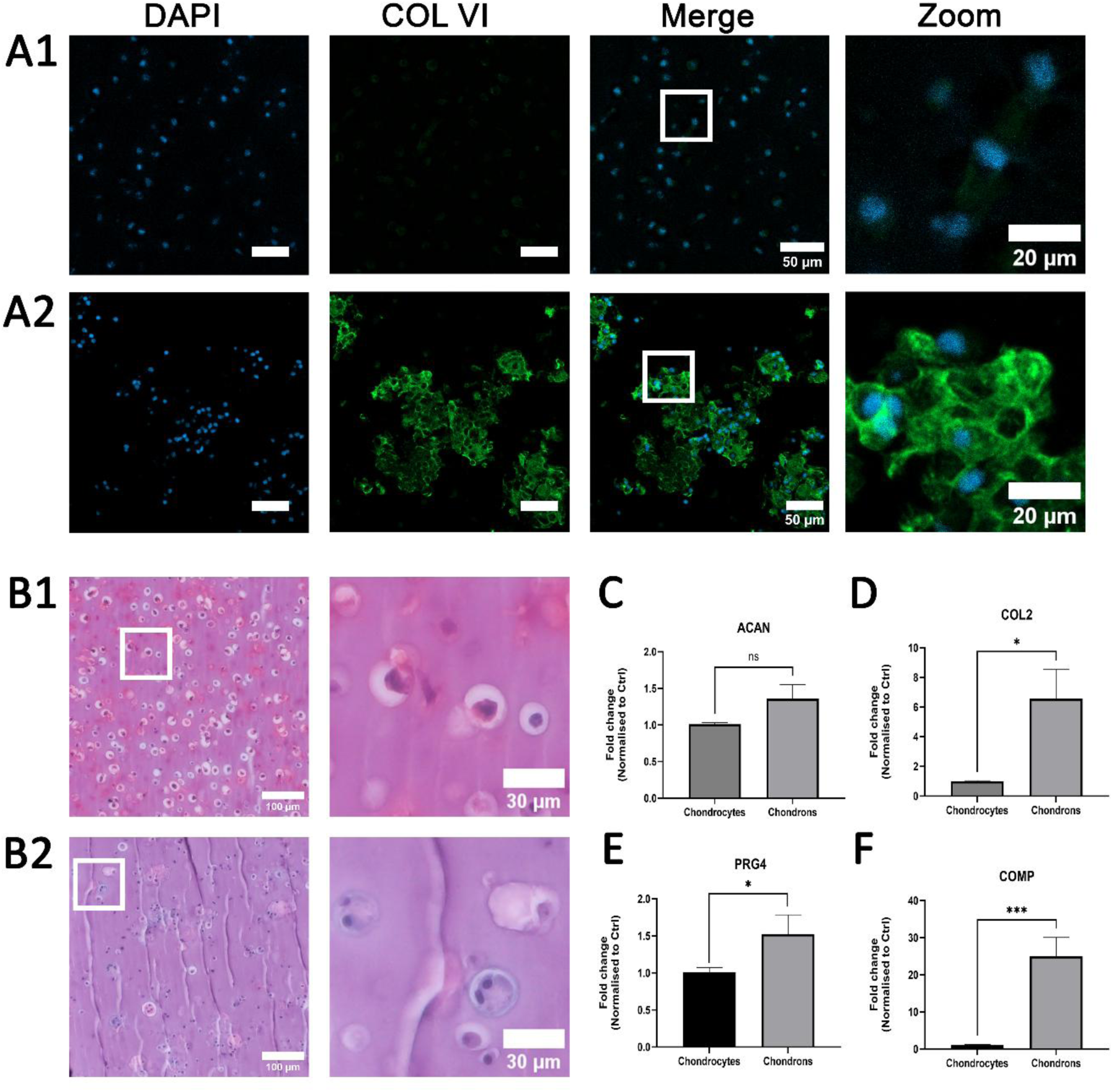
Reconstituted chondrons maintain pericellular matrix deposition and cartilage matrix synthesis in 3D GelMA culture. (A) Representative confocal immunofluorescence images of collagen type VI (COL VI, green) and nuclei (DAPI, blue) and (B) representative Safranin-O/Fast green staining in untreated and unloaded isolated chondrocytes (A1, B1) and reconstituted chondrons (A2, B2) following 7 days of culture in GelMA hydrogels. (C–F) Relative gene expression of ACAN, COL2, PRG4, and COMP in chondrocytes and reconstituted chondrons. Scale bars = 50 μm (IF) or 100 μm (bright field) for original magnification and 20μm (IF) or 30 μm (bright field) for boxed images. Bars represent mean ± SEM. N=3 donors while n=9 experimental replicates. Statistical analysis was performed using paired Student’s t-test. *P<0.05, **P<0.01, and ***P<0.001.

### 3.3 PCM reconstitution attenuates inflammatory, neuroinflammatory and pro-angiogenic gene responses under static conditions

Having confirmed that reconstituted chondrons retained a COL VI-positive PCM in 3D GelMA culture, we next examined whether PCM reconstitution altered inflammatory gene responses under static conditions. Chondrocyte and chondron GelMA constructs were cultured in the absence or presence of 1 ng/mL IL-1β for 7 days, and expression of inflammatory and catabolic genes was assessed together with NGF, a key mediator of neuro-inflammatory signaling and pain sensitization, and VEGF, a major regulator of angiogenesis during osteoarthritis. (N = 3 donors, n = 9 experimental replicates).

IL-1β induced a robust inflammatory response in chondrocytes, resulting in significant upregulation of the inflammatory mediators NOS2 (P<0.001, Fig. 3A), COX2 (P<0.001, Fig. 3B), IL-6 (P<0.001, Fig. 3C), and IL-8 (P<0.001, Fig. 3D) compared with untreated controls. In contrast, PCM-reconstituted chondrons exhibited a markedly attenuated response to IL-1β. Expression of NOS2, COX2, IL-6, and IL-8 was significantly lower than in IL-1β-treated chondrocytes (all P<0.001, Fig. 3A–D), although a significant induction of IL-8 was still observed following IL-1β stimulation in chondrons (P<0.05, Fig. 3D).

**Figure 3.**
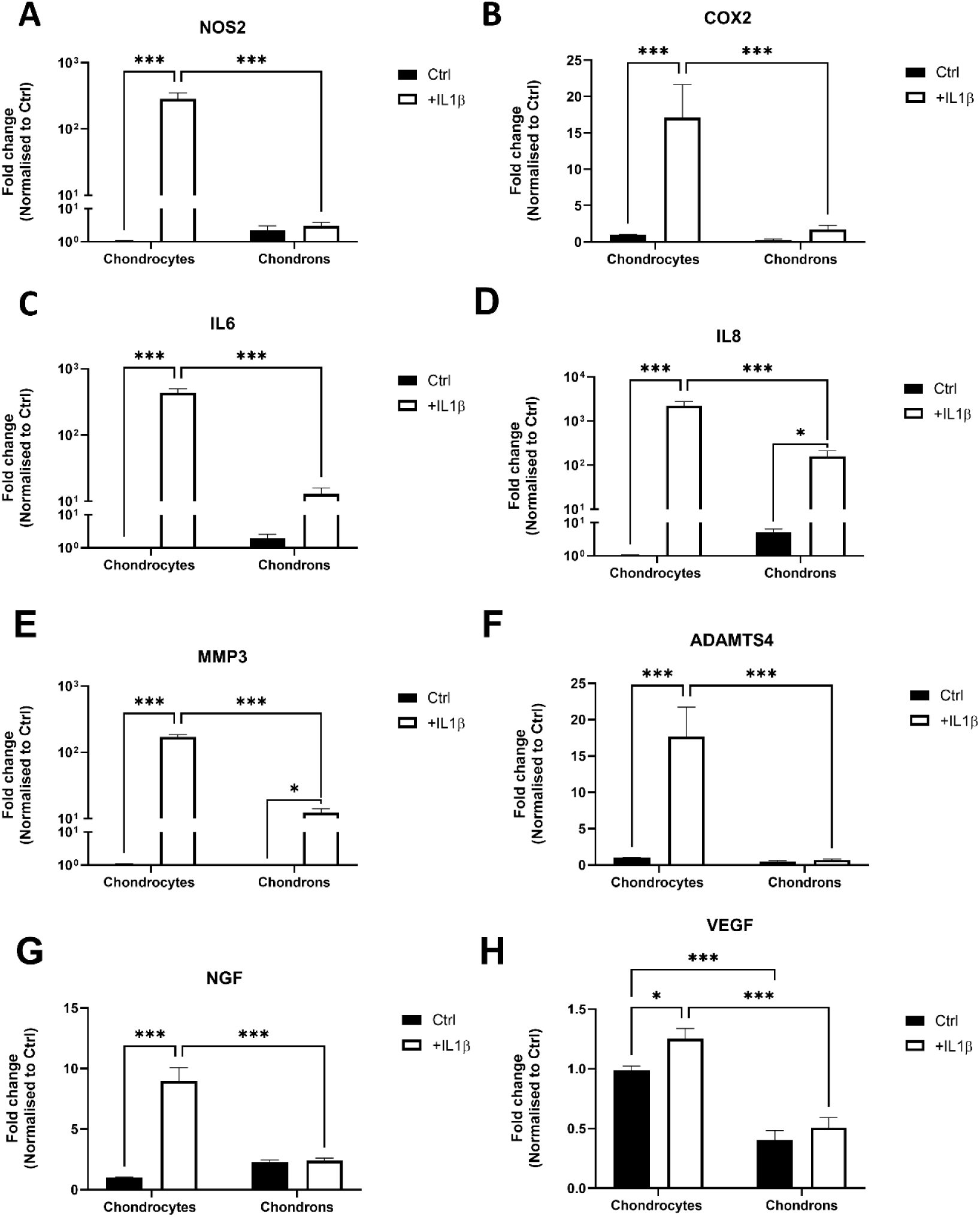
PCM reconstitution attenuates inflammatory gene responses under static conditions. Chondrocytes and chondrons were cultured under static conditions in the absence or presence of 1 ng/mL IL-1β for 7 days. Relative gene expression of NOS2 (A), COX2 (B), IL-6 (C), IL-8 (D), MMP3 (E), ADAMTS4 (F), NGF (G), and VEGF (H), normalized to the chondrocyte Ctrl group. Bars represent mean ± SEM. Statistical differences based on Two-way analysis of variance (ANOVA) with Bonferroni’s post hoc test. N=3 donors while n=9 experimental replicates Statistically significant differences are indicated as *P<0.05, **P<0.01, and ***P<0.001.

A similar pattern was observed for the catabolic markers MMP3 and ADAMTS4. IL-1β treatment significantly increased expression of both genes in chondrocytes (both P<0.001, Fig. 3E,F), whereas this response was substantially suppressed in chondrons, with significantly lower expression than IL-1β-treated chondrocytes (both P < 0.001, Fig. 3E, F). While MMP3 remained modestly inducible by IL-1β in chondrons (P<0.05, Fig. 3E), ADAMTS4 expression showed little response to inflammatory stimulation (Fig. 3F).

The neurotrophic marker NGF was also significantly upregulated by IL-1β in chondrocytes (P<0.001, Fig. 3G), whereas no significant induction was detected in chondrons, resulting in significantly lower NGF expression compared with IL-1β-treated chondrocytes (P<0.001, Fig. 3G). In contrast, VEGF exhibited a distinct expression pattern. Basal VEGF expression was significantly lower in chondrons than chondrocytes (P<0.001, Fig. 3H), and IL-1β produced only a modest increase in chondrocytes (P<0.05, Fig. 3H), while expression remained significantly lower in chondrons following IL-1β stimulation compared with chondrocytes (P<0.001, Fig. 3H).

Together, these findings demonstrate that PCM reconstitution broadly attenuates IL-1β-induced inflammatory, catabolic, neurotrophic and proangiogenic gene responses under static culture conditions.

### 3.4 PCM reconstitution attenuates inflammatory mediator release under static conditions

We next determined whether the attenuated inflammatory gene expression observed in chondrons was reflected at the protein level. Chondrocytes and chondrons were cultured under static conditions in the absence or presence of IL-1β, and release of IL-6, IL-8 and NGF was quantified by ELISA (N = 3 donors, n = 9 experimental replicates). DNA quantification revealed no significant differences in cell content between experimental groups (Supplementary Fig. S1); therefore, protein concentrations were presented as absolute values.

Consistent with the gene expression data, IL-1β induced a marked increase in IL-6 release from chondrocytes (P<0.001, Fig. 4A), whereas IL-6 release remained minimal in chondrons and was significantly lower than in IL-1β-treated chondrocytes (P<0.001, Fig. 4A). IL-8 release was also significantly increased following IL-1β stimulation in chondrocytes (P<0.001, Fig. 4B). Although IL-8 was constitutively secreted by both cell types under Ctrl conditions, basal IL-8 release was significantly lower in chondrons (P<0.05), and the response to IL-1β was markedly attenuated compared with chondrocytes (P<0.001, Fig. 4B).

**Figure 4.**
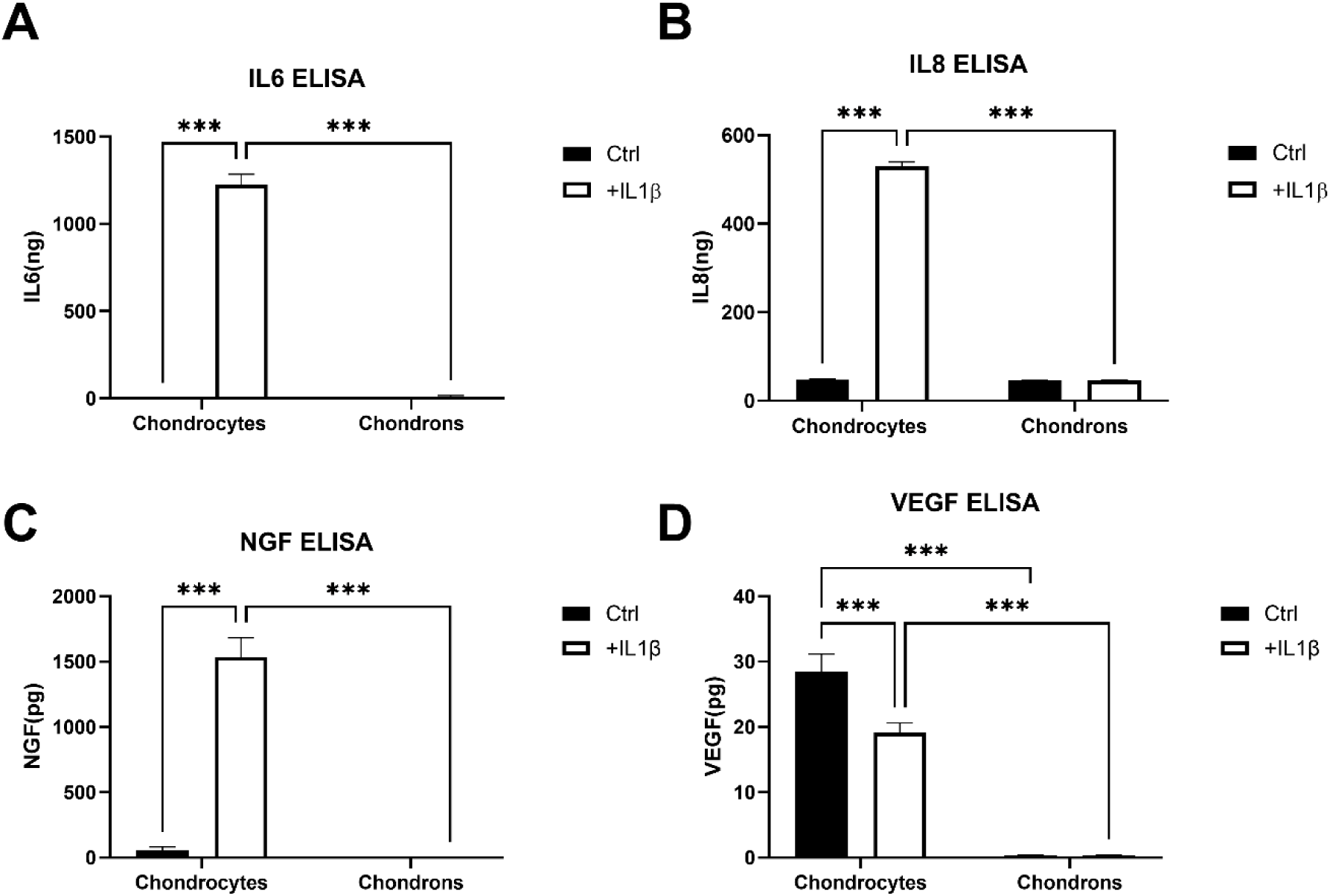
PCM reconstitution attenuates inflammatory mediator release under static conditions. Chondrocytes and chondrons were cultured under static conditions in the absence or presence of IL-1β. Release of IL-6 (A) and IL-8 (B) from 0-48h, NGF (C), and VEGF (D) release from 0-24h was quantified by ELISA. DNA content did not differ significantly between groups and is shown in Supplementary Fig. S1; protein concentrations are therefore presented as absolute values. Bars represent mean ± SEM (N = 3 independent donors, n = 9 experimental replicates). Statistical differences were assessed by two-way analysis of variance (ANOVA) with Bonferroni’s post hoc test. Statistically significant differences are indicated as *P<0.05, **P<0.01, and ***P<0.001.

**Figure 5.**
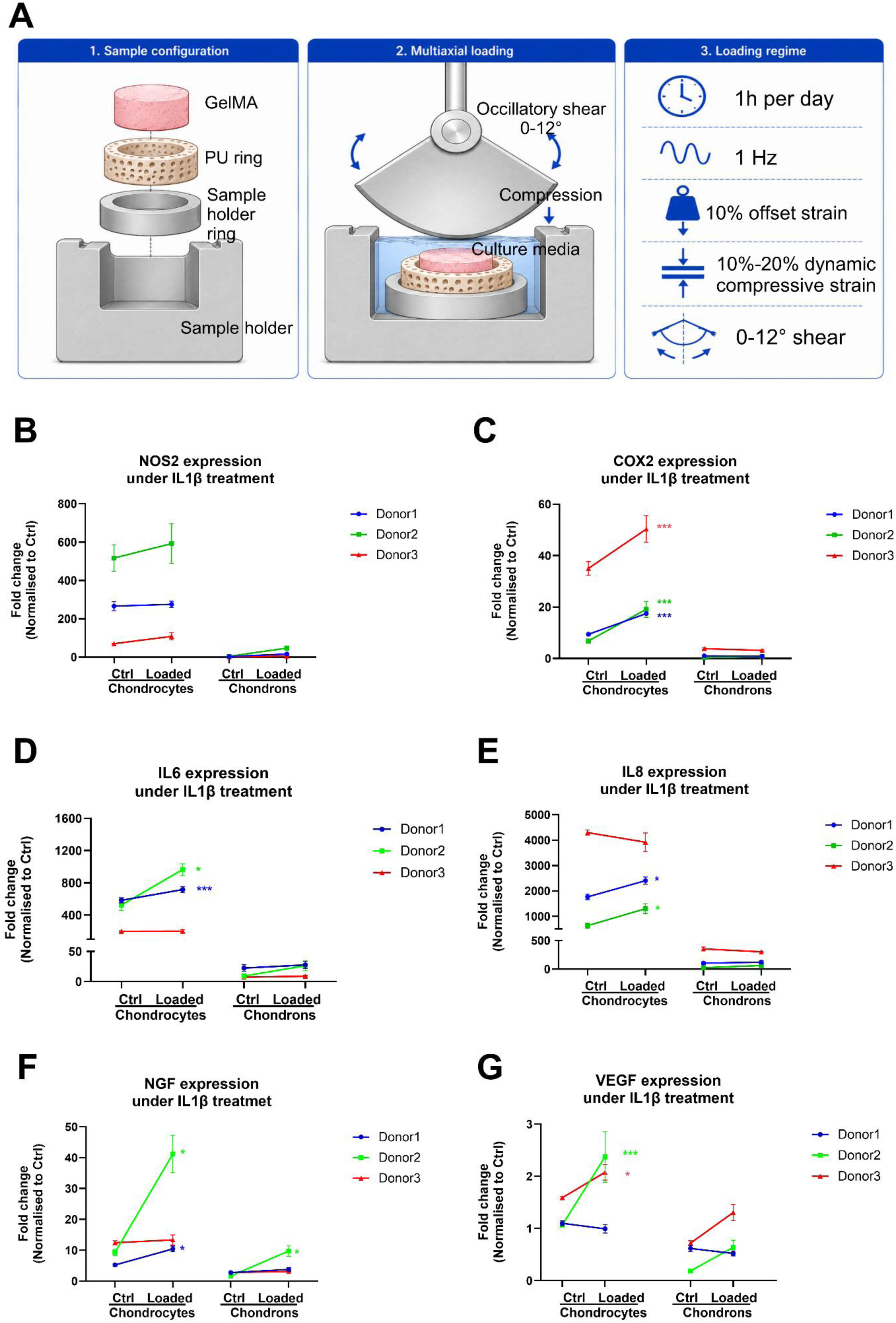
PCM buffers inflammation-induced mechanobiological responses under loading at the gene expression level. (A) Schematic of the bioreactor setup, sample configuration and multiaxial loading regime. GelMA constructs were placed within a porous polyurethane ring in the sample holder and subjected to combined compression and oscillatory shear for 1 h per day at 1 Hz, comprising 10% offset static strain, 10–20% dynamic compressive strain, and 0–12° oscillatory shear. Chondrocytes and chondrons were cultured under Ctrl or Loaded conditions in the absence or presence of IL-1β. Relative expression of NOS2 (B), COX2 (C), IL-6 (D), IL-8 (E), NGF (F), and VEGF (G) was normalized to the untreated chondrocyte Ctrl group. For clarity, only the IL-1β-treated groups are shown; complete gene-expression data are provided in Supplementary Fig. S2&3. Each coloured line represents one independent donor (N = 3, n=3 experimental replicates for each donor), highlighting donor-specific mechanobiological responses. Statistical significance indicates the effect of mechanical loading (Ctrl vs Loaded) within each donor. Statistical differences were assessed by two-way analysis of variance (ANOVA) with Bonferroni’s post hoc test. *P<0.05, **P<0.01, and ***P<0.001.

Similarly, IL-1β markedly increased NGF release by chondrocytes (P<0.001, Fig. 4C), whereas NGF remained almost undetectable in chondron condition media under both Ctrl and inflammatory conditions, resulting in significantly lower NGF release than IL-1β-treated chondrocytes (P<0.001, Fig. 4C). In contrast to the inflammatory mediators, VEGF was constitutively secreted by chondrocytes under basal conditions and was significantly reduced following IL-1β stimulation (P<0.001, Fig. 4D). VEGF release remained almost undetectable in chondrons irrespective of inflammatory stimulation and was significantly lower than that of chondrocytes under both Ctrl and IL-1β-treated conditions (both P<0.001, Fig. 4D).

Together, these findings demonstrate that the attenuated inflammatory transcriptional responses observed in PCM-reconstituted chondrons were accompanied by a corresponding reduction in inflammatory, neuro-inflammatory and angiogenic mediator release under static culture conditions.

### 3.5 PCM reconstitution buffers inflammation-induced mechanobiological responses under multiaxial loading

Having established that PCM reconstitution attenuates inflammatory responses under static conditions, we next investigated whether PCM also modulates multiaxial loading regulated inflammatory amplification. Chondrocytes are sensitive to their mechanical microenvironment[31]; therefore we have introduced a multiaxial bioreactor to mimic the physiological mechanical loading in joints (Figure 5A). Chondrocytes and chondrons were cultured under Ctrl (static) or Loaded (multiaxial loading, Figure 5A) conditions in the absence or presence of IL-1β for 7 days. As negligible loading-induced responses were observed in the absence of IL-1β (Supplementary Fig. S2&3), only the IL-1β-treated groups are presented in Fig. 5. Gene expression from each donor is shown individually (N = 3 donors, n=3 experimental replicates for each donor) to illustrate donor-specific mechanobiological responses.

Mechanical responsiveness varied between donors, with differences in both the magnitude and pattern of loading-induced responses. Nevertheless, chondrocytes consistently exhibited greater mechanosensitivity than chondrons across all donors. Among the inflammatory markers, COX2 displayed the most consistent mechanosensitive response, with loading significantly increasing expression in all three chondrocyte donors (all P<0.001, Fig. 5C). In contrast, loading-induced amplification of IL-6 and IL-8 was donor-dependent. IL-6 expression was significantly increased by loading in Donor 1 (P<0.001) and Donor 2 (P< 0.05), whereas Donor 3 showed no detectable response (Fig. 5D). Similarly, IL-8 expression was significantly increased in Donors 1 and 2 (both P<0.05), while remaining unchanged in Donor 3 (Fig. 5E). NOS2 expression remained highly elevated under inflammatory conditions in all donors but was not further enhanced by mechanical loading (Fig. 5B). In contrast to chondrocytes, expression of all four inflammatory genes remained low in chondrons, with little or no loading-induced amplification observed across donors (Fig. 5B–E).

The neuro-inflammatory mediator NGF also demonstrated donor-dependent responses under mechanical loading. Loading significantly increased NGF expression in Donors 1 and 2 (both P<0.05), whereas no significant response was observed in Donor 3 (Fig. 5F). Despite this variability, NGF expression remained consistently low in chondrons under both Ctrl and Loaded conditions. VEGF exhibited the greatest inter-donor variability, with significant loading-induced increases observed in Donors 2 (P<0.001) and 3 (P<0.05), but not in Donor 1 (Fig. 5G). In comparison, only minimal changes in VEGF expression were detected in chondrons following loading.

The matrix deposition in +/-IL-1β and +/-mechanical loading groups did not show major difference, confirmed by COLVI immunofluorescence (Supplementary Figure S4) and Safranin O/Fast Green staining (Supplementary Figure S5). Together, these findings demonstrate that although the magnitude of loading-induced responses varies substantially among donors, PCM-reconstituted chondrons consistently exhibit reduced mechanosensitivity, indicating that the regenerated pericellular matrix buffers inflammatory mechanotransduction and limits the amplification of inflammatory and neuro-inflammatory signaling during mechanical loading.

### 3.6 PCM reconstitution maintains a reduced inflammatory secretome under mechanical loading

We next examined whether the transcriptional responses to mechanical loading were reflected at the protein level. Chondrocytes and chondrons were cultured under Ctrl or Loaded conditions in the absence or presence of IL-1β. Similarly, as mechanical loading produced minimal changes in protein release in the absence of inflammatory stimulation, only the IL-1β-treated groups are presented in the main figure for clarity, while the complete dataset is provided in Supplementary Fig. S1. Release of IL-6, IL-8, and NGF were quantified by ELISA (N = 3 donors, n = 9 experimental replicates).

Consistent with the transcriptional data, release of all three mediators remained markedly lower in chondrons than in chondrocytes under both Ctrl and Loaded conditions. IL-6 release remained significantly lower in chondrons than in chondrocytes under both mechanical conditions (both P < 0.001, Fig. 6A). Although mechanical loading produced a modest increase in IL-6 release in chondrocytes, these changes did not reach statistical significance (P>0.05). Similarly, IL-8 release was significantly reduced in chondrons compared with chondrocytes under both Ctrl and Loaded conditions (both P < 0.001, Fig. 6B). Mechanical loading likewise increased IL-8 release, reaching statistical significance only in chondrocytes (P<0.01), whereas the increase observed in chondrons did not reach significance.

**Figure 6.**
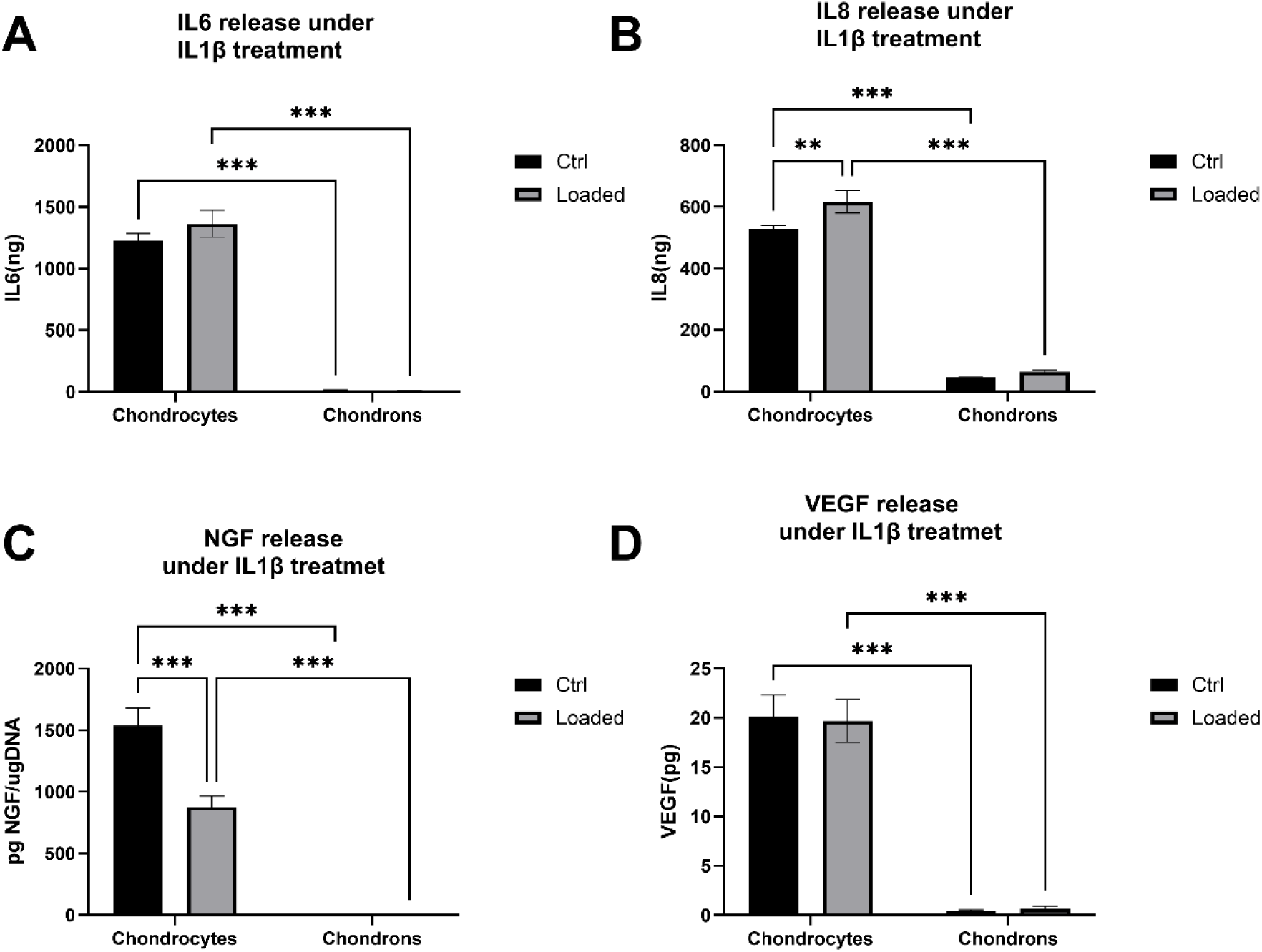
PCM maintains a reduced inflammatory secretome under mechanical loading. Chondrocytes and chondrons were cultured under Ctrl or Loaded conditions in the absence or presence of IL-1β. Release of IL-6 (A) and IL-8 (B) from 0-48h, NGF (C), and VEGF (D) release from 0-24h was quantified by ELISA. DNA content did not differ significantly between groups and is shown in Supplementary Fig. S1; protein concentrations are therefore presented as absolute values. For clarity, only the IL-1β-treated groups are shown; complete ELISA data, including untreated controls, are provided in Supplementary Fig. S1. Bars represent mean ± SEM (N = 3 independent donors, n = 9 experimental replicates). Statistical differences were assessed by two-way analysis of variance (ANOVA) with Bonferroni’s post hoc test. Statistically significant differences are indicated as *P<0.05, **P<0.01, and ***P<0.001.

**Figure 7.**
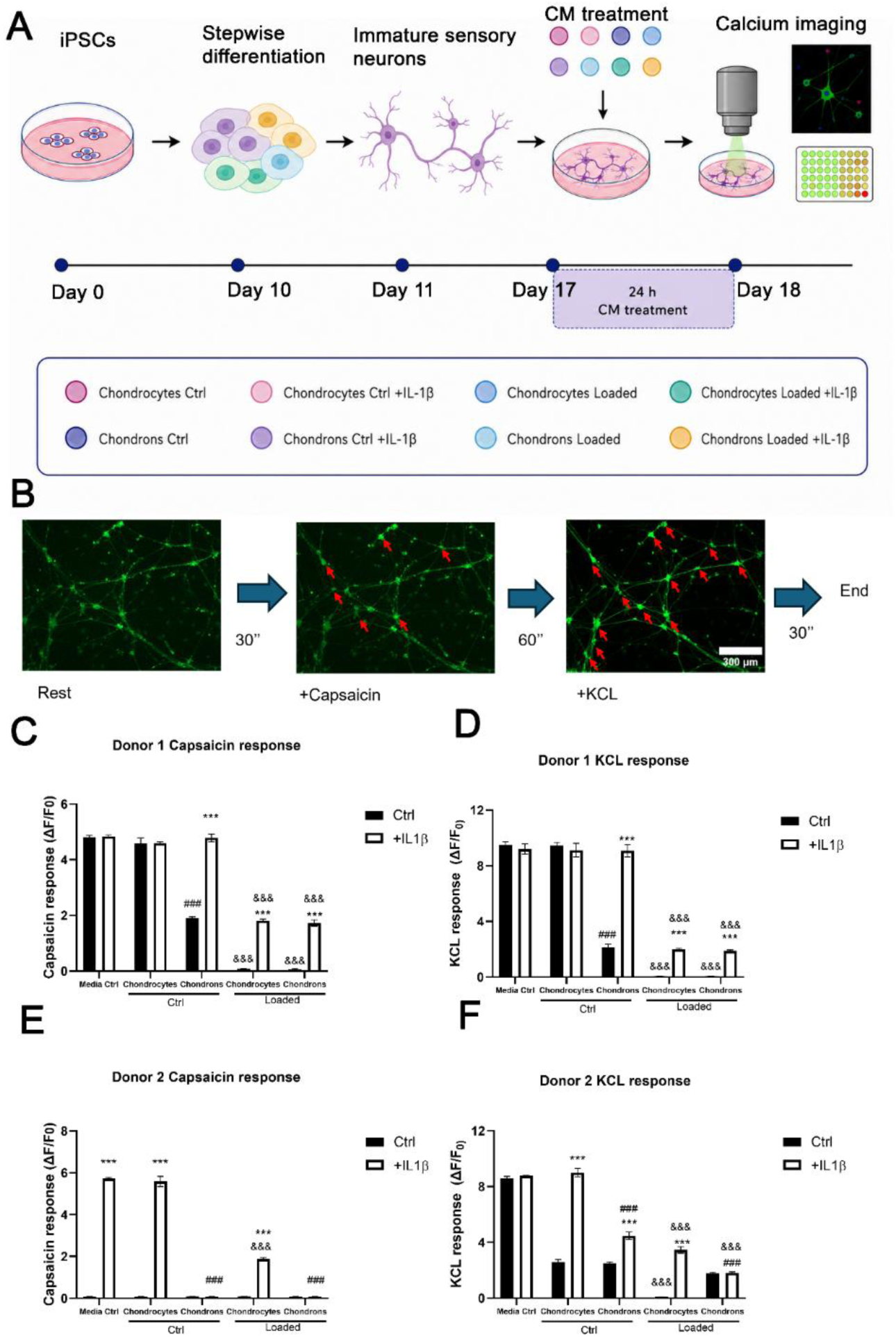
Conditioned media modulate sensory-neuron calcium responses. (A) Schematic overview of the iPSC-derived sensory neuron (iPSC-SN) experimental workflow. Mature iPSC-SNs were generated by stepwise differentiation and exposed to conditioned media (CM) for 24 h prior to functional assessment. (B) Schematic representation of the calcium imaging assay showing baseline recording followed by capsaicin and KCl stimulation. (C, D) Capsaicin-evoked and KCl-evoked calcium peak responses following treatment with conditioned media from cartilage donor 1. (E, F) Capsaicin-evoked and KCl-evoked peak calcium responses following treatment with conditioned media from cartilage donor 2. Statistical significance indicates the effect of IL-1β treatment (***), differences between chondrocyte-and chondron-derived conditioned media (###), and the effect of mechanical loading (&&&) within the corresponding groups. Bars represent mean ± SEM. N=2 donors while n=6 experimental replicates. Statistical differences based on Two-way analysis of variance (ANOVA) with Bonferroni’s post hoc test.

NGF release displayed a distinct response. Although NGF release remained significantly lower in chondrons than chondrocytes under both Ctrl and Loaded conditions (both P<0.001, Fig. 6C), mechanical loading significantly reduced NGF release in chondrocytes (P<0.001), whereas no loading-induced change was observed in chondrons (Fig. 6C). VEGF release exhibited a similar pattern to that observed under static conditions. VEGF remained almost undetectable in chondrons irrespective of mechanical loading and was significantly lower than in chondrocytes under both Ctrl and Loaded conditions (both P<0.001, Fig. 6D). Mechanical loading did not significantly alter VEGF release within either cell type (P>0.05).

Together, these findings demonstrate that, while mechanical loading tended to enhance the inflammatory secretory profile of IL-1β-treated chondrocytes, PCM-reconstituted chondrons maintained consistently low release of inflammatory, neuro-inflammatory and angiogenic mediators irrespective of loading. These findings suggest that PCM reconstitution not only suppresses inflammatory mediator release but also buffers loading-induced amplification of the inflammatory secretome.

### 3.7 Conditioned media from reconstituted chondrons modulate sensory-neuron calcium responses

As an exploratory assessment of cartilage–sensory neuron crosstalk, we examined whether conditioned media (CM) generated by chondrocytes and reconstituted chondrons under inflammatory and mechanical conditions altered the functional responsiveness of iPSC-derived sensory neurons (iPSC-SNs). iPSC-derived sensory neurons (iPSC-SNs) were generated using a stepwise differentiation protocol (see section 2.7 and Supplementary Figure S6 for details) and treated for 24h with conditioned media (CM) collected 0-24h from the eight culture conditions (chondrocytes or chondrons, ±IL-1β, Ctrl or Loaded), plus a chondropermissive media control (Media Ctrl) and a chondropermissive media + 1ng/mL IL-1β group (Media Ctrl +IL-1β), prior to calcium imaging (Fig. 7A, B). Conditioned media from two independent donors were evaluated in separate iPSC-SN experiments and presented individually.

Although the magnitude and IL-1β dependence of neuronal responses differed between the two experiments, several consistent patterns were observed. Under static conditions (Ctrl), chondron-derived CM generally produced lower capsaicin-and KCl-evoked calcium responses than the corresponding chondrocyte-derived CM, with significant differences observed in both experiments (P < 0.001; Fig. 7C–F). Mechanical loading further reduced evoked calcium responses in matched chondrocyte and chondron CM conditions (P < 0.001, Fig. 7C–F), with loaded chondron-derived CM consistently associated with low neuronal responses. The effect of IL-1β differed between the two experiments. In Donor 1, IL-1β had little effect on the Media Ctrl and Ctrl chondrocyte-CM groups but increased capsaicin-and KCl-evoked responses in the chondron-CM groups (Fig. 7C-D). In Donor 2, IL-1β markedly increased responses in the Media Ctrl and Ctrl chondrocyte-CM groups, whereas its effect on chondron-derived CM was weaker and depended on the stimulus and loading condition (Figure 7E-F).

Across both experiments, capsaicin-and KCl-evoked responses generally changed in parallel, suggesting modulation of overall neuronal calcium responsiveness rather than a selective effect on TRPV1-mediated activation. These functional changes were not accompanied by detectable differences in neuronal morphology, the proportion of CGRP-positive sensory neurons, or neuronal viability (Supplementary Figs. S7 and S8), while representative calcium traces are provided in Supplementary Fig. S9. Together, these exploratory findings demonstrate that CM modulates sensory-neuron calcium responsiveness, with chondron-derived CM producing lower capsaicin-and KCl-evoked responses than corresponding chondrocyte-derived CM in key comparisons across both independent experiments.

## 4. Discussion

In this study, we investigated whether reconstruction of the pericellular microenvironment lost during chondrocyte isolation could generate a functionally distinct chondron-like cellular state. Alginate preconditioning promoted formation of a COL VI-rich PCM that was retained following transfer to 3D GelMA culture and was accompanied by increased cartilage matrix deposition (Figs. 1 and 2). As hypothesized, reconstituted chondrons showed broadly attenuated responses to IL-1β, encompassing inflammatory, catabolic, neurotrophic, and angiogenic markers at both gene and protein levels (Figs. 3 and 4). Under combined inflammatory and mechanical challenge, however, the response was more complex than a uniform effect of loading: chondrocytes exhibited marker-and donor-dependent transcriptional responses, whereas reconstituted chondrons generally maintained substantially lower inflammatory activity and consistently low mediator secretion (Figs. 5 and 6). Notably, loading-induced transcriptional changes were not always reflected by corresponding changes in secreted protein concentrations, indicating additional regulation between gene expression and protein release (Figs. 5 and 6). Exploratory conditioned-medium experiments further showed that chondron-derived media were associated with lower capsaicin-and KCl-evoked sensory-neuron calcium responses in key comparisons across both independent experiments, although the magnitude and inflammatory dependence of these effects varied (Fig. 7). Collectively, these findings establish that the reconstituted chondron state is functionally distinct from isolated chondrocytes and is associated with greater resilience to inflammatory and mechanical challenges.

### 4.1 PCM reconstitution and inflammatory protection

Previous studies have shown that chondrocytes can regenerate components of the PCM during three-dimensional culture[32–36]. Consistent with these reports, alginate preconditioning in the present study generated chondron-like structures surrounded by a distinct COL VI-positive matrix (Fig. 1). Importantly, this pericellular matrix was retained after transfer into GelMA, indicating that the reconstructed phenotype was maintained beyond the alginate preconditioning phase (Fig. 2). Reconstituted chondrons also showed greater cartilage matrix deposition and increased COL2 and COMP expression, whereas ACAN and PRG4 were not significantly altered (Fig. 2). Thus, PCM reconstitution was associated with selective enhancement of cartilage matrix-related features rather than a uniform increase across all chondrogenic markers[32].

The more striking functional difference emerged following inflammatory stimulation. IL-1β induced robust inflammatory, catabolic and neurotrophic responses in isolated chondrocytes, whereas these responses were broadly attenuated in reconstituted chondrons, including NOS2, COX2, IL6, IL8, MMP3, ADAMTS4 and NGF expression (Fig. 3). This difference was also evident at the protein level, with markedly lower IL6, IL8 and NGF secretion from chondrons and consistently low VEGF release (Fig. 4). The breadth of these differences suggests that reconstruction of the chondron-like microenvironment is associated with a generally less inflammatory and catabolic cellular state, rather than regulation of a single pathway. Such an effect is compatible with established roles of the PCM in controlling the local presentation of soluble factors and cell–matrix interactions[8, 9, 16, 37], although these mechanisms were not directly examined here. Importantly, the reconstitution procedure also altered cell organization and promoted broader matrix deposition; therefore, the present study cannot attribute the observed functional differences exclusively to COL VI or to the PCM independently of cell aggregation and other cell–matrix interactions. Future studies using PCM-specific perturbations will be required to distinguish these contributions.

### 4.2 The PCM and mechanically regulated inflammatory response

The response to mechanical loading revealed an important distinction between isolated chondrocytes and reconstituted chondrons. In the absence of IL-1β, loading produced relatively limited changes, whereas under inflammatory stimulation it further modified the expression of several inflammatory and neurotrophic genes in chondrocytes (Fig. 5 and Supplementary Fig. S2). This effect was not uniform: COX2 showed the most consistent loading-associated increase across donors, while IL6, IL8, NGF and VEGF displayed greater inter-donor variability, and NOS2 showed little additional response to loading. These findings highlight that inflammatory mechanobiology is both marker-and donor-dependent and suggest that mechanical loading interacts with the existing inflammatory state rather than acting as a uniform pro-inflammatory stimulus. In contrast, reconstituted chondrons generally maintained substantially lower expression across these markers and showed less pronounced loading-associated changes (Fig. 5), indicating that the chondron-like state was associated with reduced sensitivity to inflammatory mechanical challenge.

Interestingly, the transcriptional responses to loading were not uniformly reproduced at the protein level (Fig. 6). IL6 and IL8 secretion showed increasing trends following loading but did not change significantly, while NGF secretion decreased significantly in loaded chondrocytes and VEGF remained largely unaffected. Thus, loading-induced changes in gene expression should not be interpreted as directly equivalent to increased mediator release. Such divergence may reflect differences in transcriptional kinetics, protein synthesis, secretion, degradation, or the timing of conditioned-medium collection. Despite these marker-specific responses in chondrocytes, reconstituted chondrons maintained consistently low or nearly undetectable secretion of IL6, IL8, NGF and VEGF under both Ctrl and Loaded conditions (Fig. 6). The stability of this secretory phenotype, despite donor-dependent transcriptional responses, represents one of the more robust findings of the present study and suggests that the reconstituted chondron state limits loading-associated inflammatory output at multiple levels.

The basis for this altered mechanobiological responsiveness was not directly investigated. However, established properties of the native PCM provide two complementary hypotheses (Fig. 8). Matrix-associated molecules such as perlecan can retain and regulate the availability of growth factors, supporting a biochemical “reservoir” function[8, 37–39] (Fig. 8A), while the compliant and highly hydrated PCM can redistribute local stress, fluid flow and mechano-osmotic cues before they reach the cell membrane like a “water balloon”[5, 40–45] (Fig. 8A). These biochemical and mechanical functions may therefore modify the cellular inputs generated by an identical tissue-level loading regime. Importantly, neither PCM mechanics nor matrix-bound mediator release was directly measured here; the proposed reservoir and mechanical-buffer models should therefore be regarded as conceptual explanations for the observed differences rather than mechanisms demonstrated by the present study (Fig. 8).

**Figure 8.**
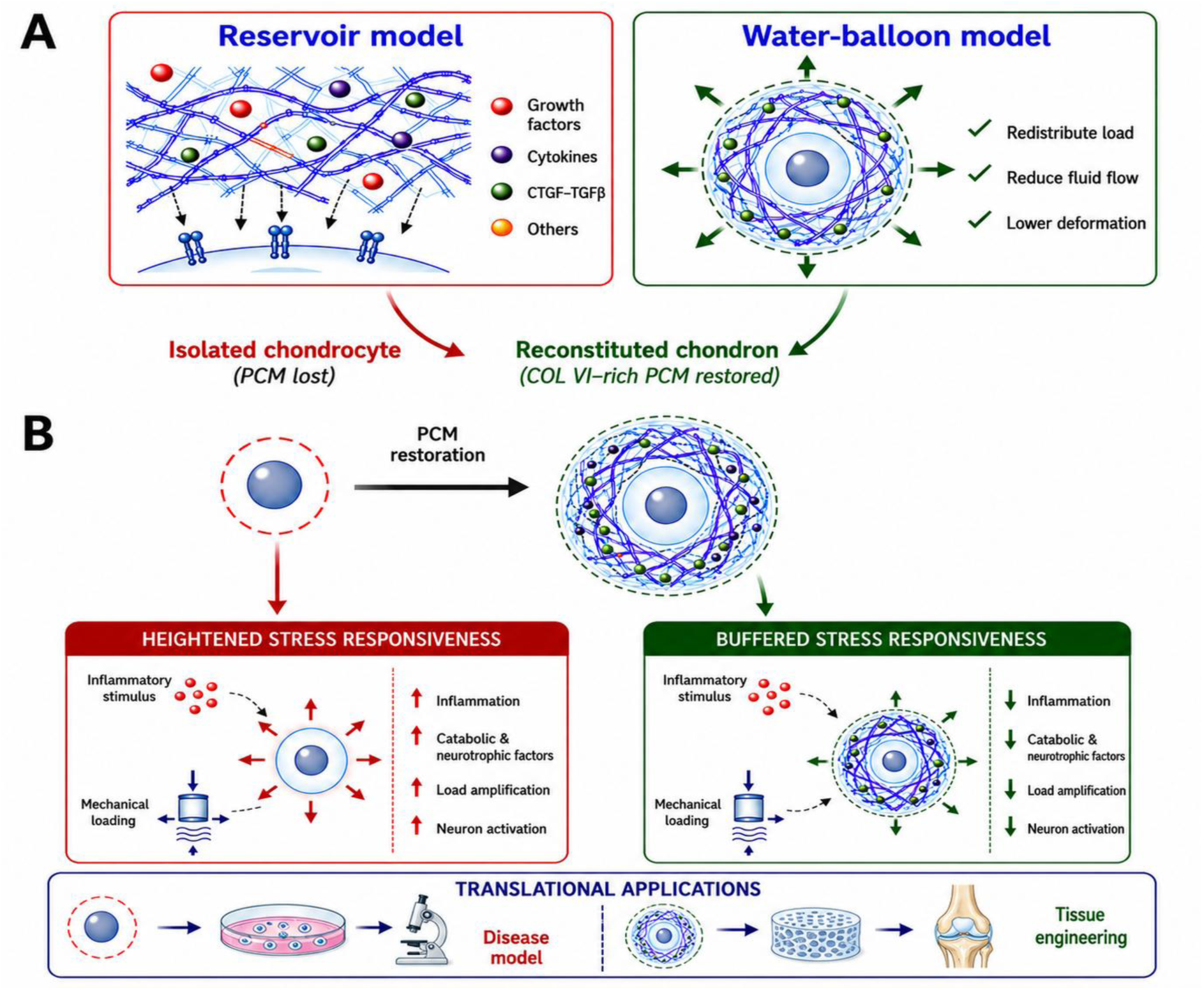
Proposed mechanisms and translational significance of PCM restoration. (A) Two complementary models are proposed to explain how a restored COL VI–rich pericellular matrix (PCM) modulates chondrocyte responses. In the “reservoir” model, the PCM retains and organizes soluble mediators, thereby regulating their local availability. In the “water-balloon” model, the hydrated PCM redistributes mechanical load and limits cellular deformation. (B) These mechanisms may account for the heightened stress responsiveness of PCM-deficient isolated chondrocytes and the buffered responses of reconstituted chondrons. Translationally, the conventional chondrocyte model may overestimate cellular sensitivity to external stimuli, whereas the reconstituted chondron model may better reproduce the native chondrocyte microenvironment for studying osteoarthritis-related responses.

### 4.3 PCM-mediated regulation of cartilage–sensory neuron crosstalk

The conditioned-medium experiments were included as an exploratory functional study to determine whether the distinct secretory profiles of isolated chondrocytes and reconstituted chondrons could translate into differences in sensory-neuron responsiveness. Across both independent experiments, chondron-derived CM was associated with lower capsaicin-and KCl-evoked calcium responses than the corresponding chondrocyte-derived CM in key comparisons, while mechanical loading further reduced neuronal responses in several conditions (Fig. 7). These findings are consistent with the markedly lower release of inflammatory and neurotrophic mediators from reconstituted chondrons (Figs. 4 and 6). NGF represents one plausible contributor, as inflammatory and mechanical stimulation can increase chondrocyte NGF production[46, 47]and NGF can enhance nociceptor responsiveness[48]. However, the present experiments do not establish a causal relationship between any individual cartilage-derived mediator and the neuronal response.

The two sensory-neuron experiments nevertheless showed notable differences in baseline and stimulus-evoked responsiveness. For example, capsaicin-and KCl-evoked responses in the media-control groups differed between experiments, and chondron-derived CM produced minimal capsaicin responses under both ±IL-1β conditions in one experiment but not to the same extent in the other (Fig. 7). One possible explanation is variability in the maturation and functional sensitivity of the iPSC-SNs themselves. Although the neurons were derived from the same cell batch using the same differentiation protocol and culture duration, their calcium responsiveness differed between independent experiments, suggesting that absolute response amplitudes should not be directly cross-referenced across experimental runs. A second source of variability may arise from donor-dependent differences in the composition of the cartilage-derived CM. Given that iPSC-SN calcium imaging is not readily suited to high-throughput testing, these observations support interpreting treatment effects primarily within each experimental run rather than across separate experiments. For future studies incorporating additional cartilage donors, a more robust design would be to test multiple donor-derived conditioned media in parallel within the same neuronal preparation, thereby allowing donor-specific effects to be compared under a common experimental background.

### 4.4 Clinical implications and translational significance

The present study demonstrates that PCM depletion and PCM restoration generate two distinct cellular phenotypes with fundamentally different responses to inflammatory, and mechanical stimuli, as well as inducing distinct neuroactivity. Throughout this study, PCM-depleted chondrocytes consistently exhibited amplified inflammatory and mechano-inflammatory responses, whereas reconstituted chondrons maintained a more homeostatic phenotype under the same experimental conditions. These findings suggest that the presence or absence of PCM is not merely a structural characteristic, but a key determinant of chondrocyte behavior and cellular resilience.

The heightened responsiveness of PCM-depleted chondrocytes may be advantageous for disease modelling. As enzymatic isolation inevitably removes the native PCM, isolated chondrocytes closely resemble the cellular state currently used in most *in vitro* cartilage studies. Rather than considering this as a limitation, we propose that isolated chondrocytes represent a stress-amplifying system, capable of maximizing inflammatory and mechano-inflammatory responses and thereby facilitating the investigation of osteoarthritis pathophysiology, mechanobiology, cartilage–sensory neuron crosstalk, and therapeutic screening (Fig. 8B).

Conversely, reconstituted chondrons consistently demonstrated enhanced resistance to inflammatory and mechanical perturbation, supporting their potential application in regenerative medicine. By restoring the native pericellular niche, chondrons appear to function as a stress-buffering system, preserving cellular homeostasis under adverse conditions. This phenotype may be particularly advantageous for cartilage tissue engineering and cell-based repair strategies, where long-term survival and functional stability are essential (Fig. 8B). Together, these findings suggest that isolated chondrocytes and reconstituted chondrons should not be viewed as competing cell models, but as complementary platforms optimized for different translational applications.

### 4.5 Limitations and future perspective

Several limitations of this study should be acknowledged. Although our findings demonstrate that PCM restoration functionally regulates inflammatory, mechanobiological, and cartilage–sensory neuron responses, the underlying molecular mechanisms remain to be fully elucidated. In particular, the specific matrix-associated mediators and mechanotransduction pathways responsible for these protective effects require further investigation. Future studies integrating proteomic analyses, direct chondrocyte–sensory neuron co-culture systems, and *in vivo* models will be important to establish the molecular basis and translational relevance of PCM restoration.

## 5. Conclusions

This study demonstrates that a collagen VI-rich pericellular matrix can be successfully reconstituted around isolated human chondrocytes and maintained in 3D culture, generating chondron-like cellular units with attenuated inflammatory activation and buffered loading-associated inflammatory responses. Exploratory conditioned-medium experiments further showed that reconstituted chondrons were generally associated with lower capsaicin-and KCl-evoked sensory-neuron calcium responses than isolated chondrocytes. Together, these findings suggest that PCM functions as a mechanochemical buffer that modulates chondrocyte responses to biochemical and mechanical challenges. Restoring the pericellular microenvironment lost during cell isolation and expansion may therefore represent a useful design principle for cartilage tissue engineering and cell-based repair.

## Declaration of generative AI and AI-assisted technologies in the manuscript preparation process

During the preparation of this work, the authors used ChatGPT (GPT-5.6 Sol) in order to polish grammar and check coherence and spelling. After using this tool, the authors reviewed and edited the content as needed and took full responsibility for the content of the published article.

## Supplementary Information

**Figure S1.**
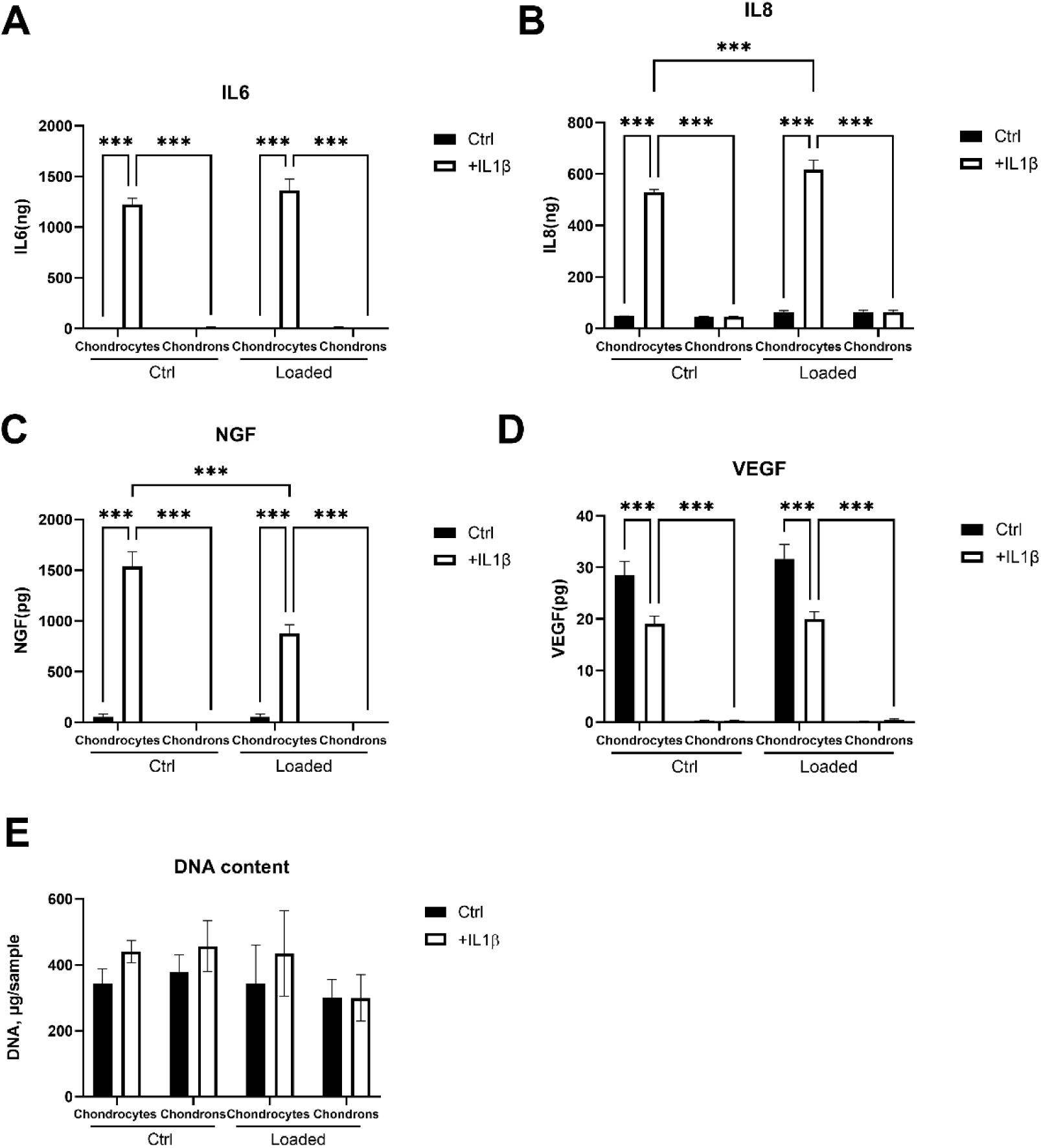
Complete ELISA quantification of inflammatory and neuroinflammatory mediators under free swelling and loaded culture conditions. 0-48h cumulative secretion of IL6 (A), IL8 (B), and 0-24h cumulative secretion of NGF (C), and VEGFA (D) by chondrocytes and reconstituted chondrons cultured under control (Ctrl) or IL-1β-stimulated (+IL-1β) conditions in the absence (Ctrl) or presence (Loaded) of mechanical loading. Absolute protein contents measured by ELISA are presented without normalization to DNA content. DNA content per construct is shown in (E), demonstrating no significant differences between experimental groups. Bars represent mean ± SEM (N = 3 independent donors, n = 9 experimental replicates). Statistical analysis was performed using two-way analysis of variance (ANOVA) with Bonferroni’s post hoc test. *P<0.05, **P< 0.01, ***P<0.001.

**Figure S2.**
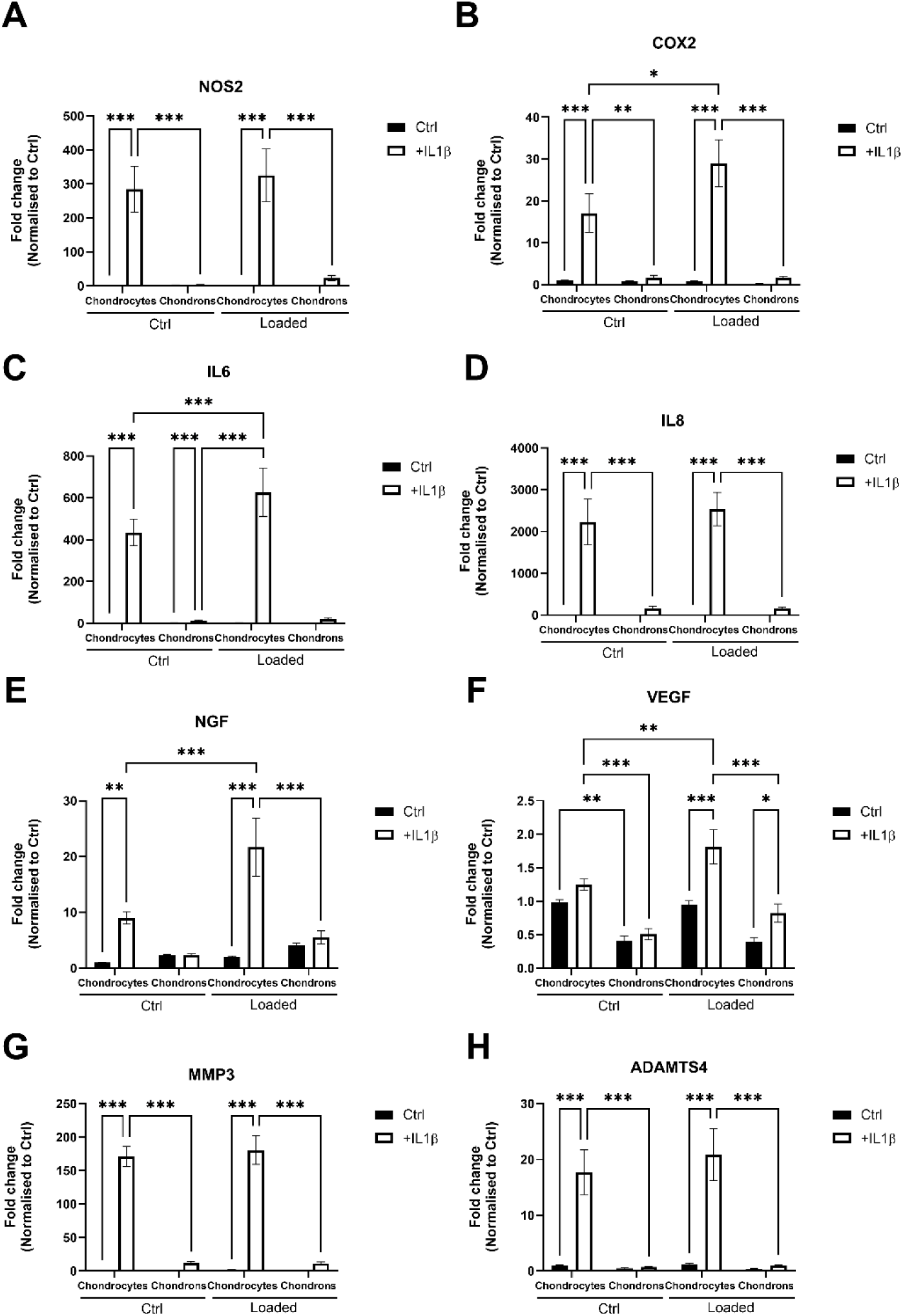
Extended inflammatory, catabolic and neurovascular-related gene expression analysis under all culture conditions. Expression of NOS2 (A), PTGS2 (COX2) (B), IL6 (C), CXCL8 (IL8) (D), NGF (E), VEGFA (F), MMP3 (G), and ADAMTS4 (H) in chondrocytes and reconstituted chondrons cultured under control (Ctrl) or IL-1β-stimulated (+IL-1β) conditions +/-mechanical loading. Gene expression was quantified by RT-qPCR and normalized to the housekeeping gene RPLP0. Data are presented as fold change relative to untreated chondrocytes (Ctrl). Bars represent mean ± SEM (N = 3 donors, n=9 experimental replicates). Statistical analysis was performed using two-way analysis of variance (ANOVA) with Bonferroni’s post hoc test. *P<0.05, **P< 0.01, ***P<0.001.

**Figure S3.**
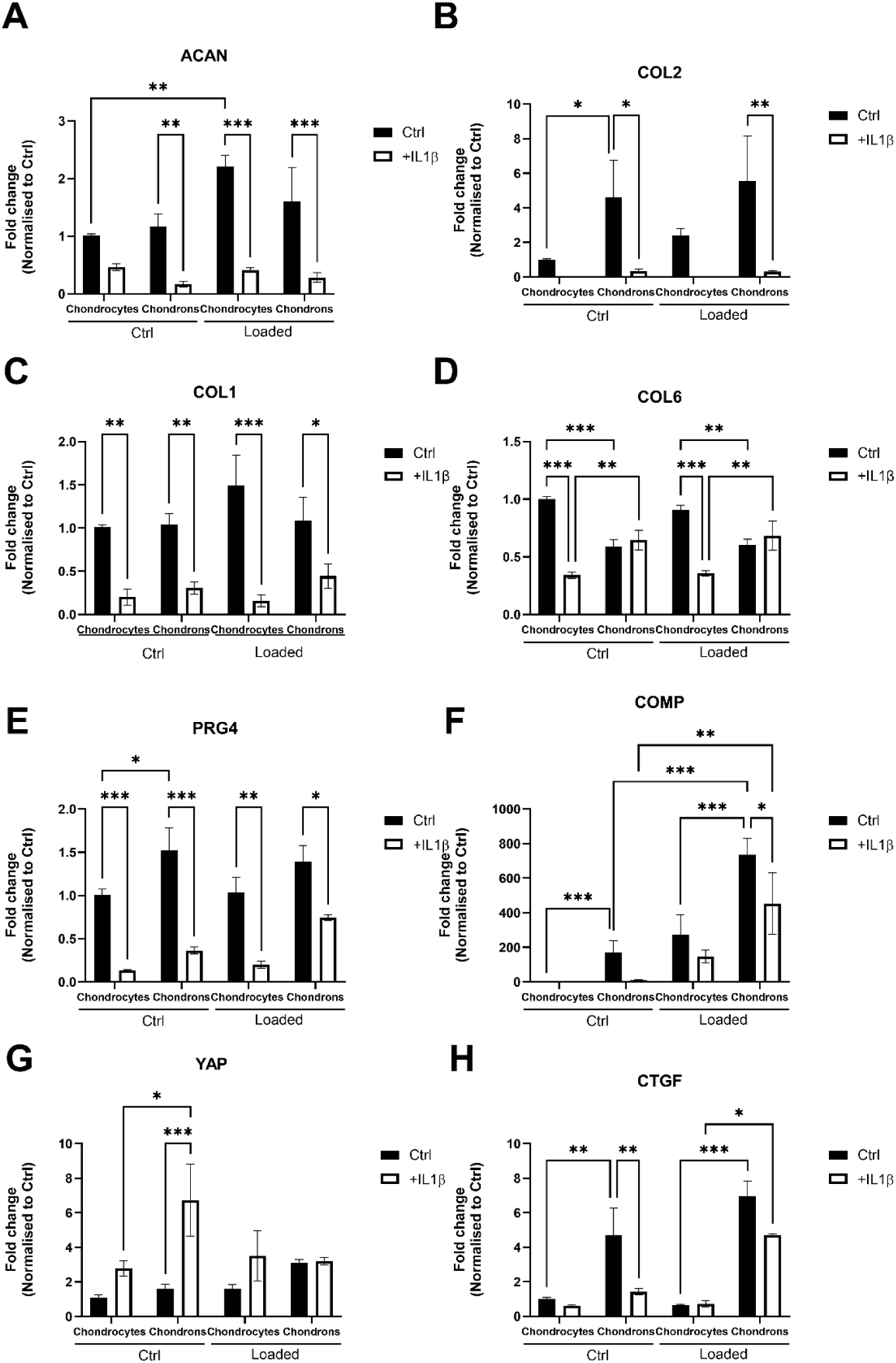
Extended analysis of cartilage matrix-associated and mechanotransduction-related gene expression under all culture conditions. Expression of ACAN (A), COL2A1 (B), COL1A1 (C), COL6A1 (D), PRG4 (E), COMP (F), YAP1 (G), and CTGF (H) in chondrocytes and reconstituted chondrons cultured under control (Ctrl) or IL-1β-stimulated (+IL-1β) conditions +/-mechanical loading. Gene expression was quantified by RT-qPCR and normalized to the housekeeping gene RPLP0. Data are presented as fold change relative to untreated chondrocytes (Ctrl). Bars represent mean ± SEM (N = 3 donors, n=9 experimental replicates). Statistical analysis was performed using two-way analysis of variance (ANOVA) with Bonferroni’s post hoc test. *P<0.05, **P< 0.01, ***P<0.001.

**Figure S4.**
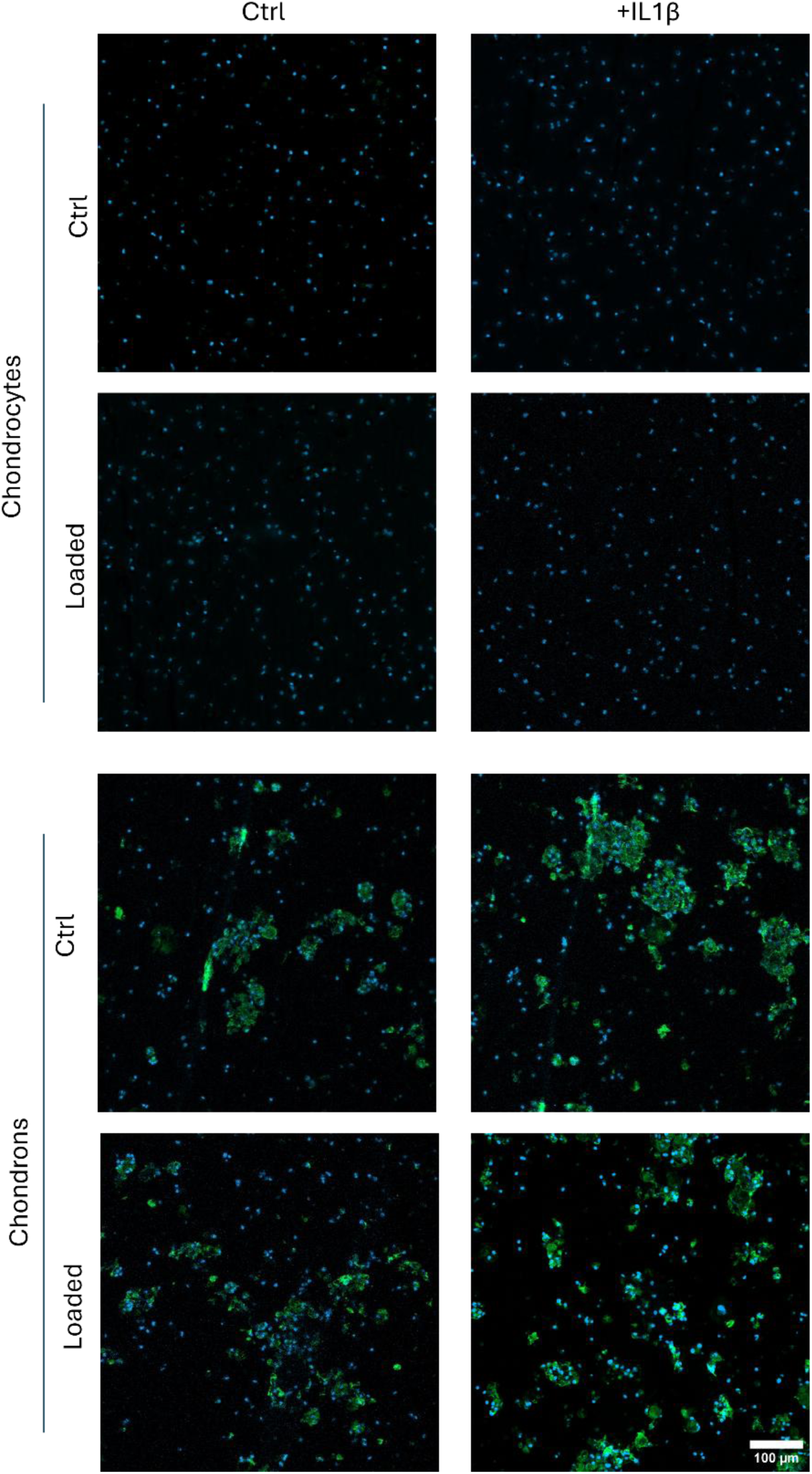
Collagen VI immunofluorescence staining of GelMA constructs under all experimental conditions. Representative immunofluorescence images showing collagen VI (green) deposition in GelMA constructs containing chondrocytes (upper section) or reconstituted chondrons (lower section) cultured under control (Ctrl, left column) or IL-1β-stimulated (+IL-1β, right column) conditions +/-mechanical loading. Cell nuclei were counterstained with DAPI (blue). Representative images from. Scale bar = 100 μm.

**Figure S5.**
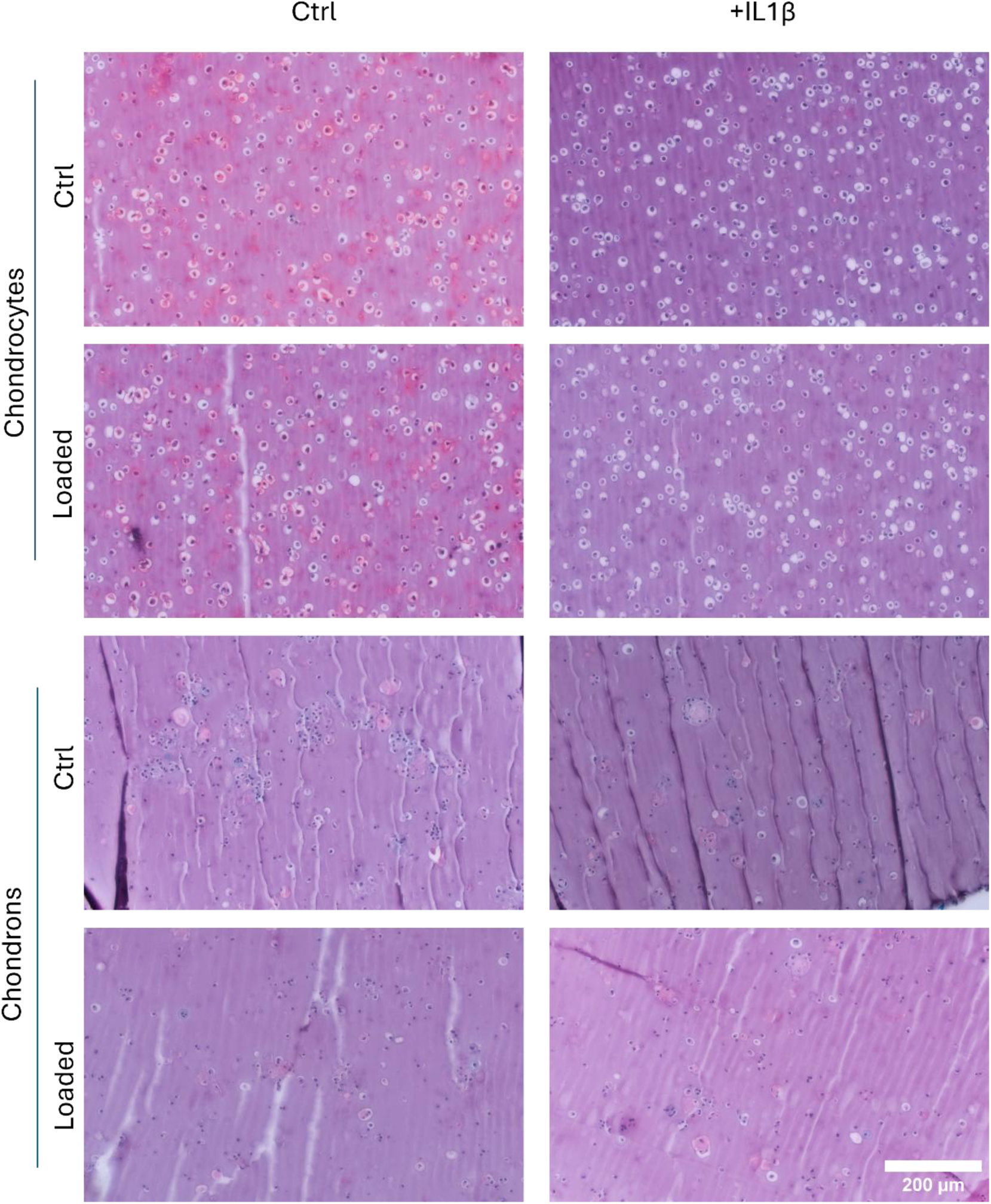
Safranin O/Fast Green staining of GelMA constructs under all experimental conditions. Representative Safranin O/Fast Green-stained sections of GelMA constructs containing chondrocytes or reconstituted chondrons cultured under control (Ctrl, left column) or IL-1β-stimulated (+IL-1β, right column) conditions in the absence (upper panels) or presence (lower panels) of mechanical loading. Safranin O stains sulfated glycosaminoglycans (red), while Fast Green counterstains the surrounding matrix (green-blue).. Scale bar = 200 μm.

**Figure S6.**
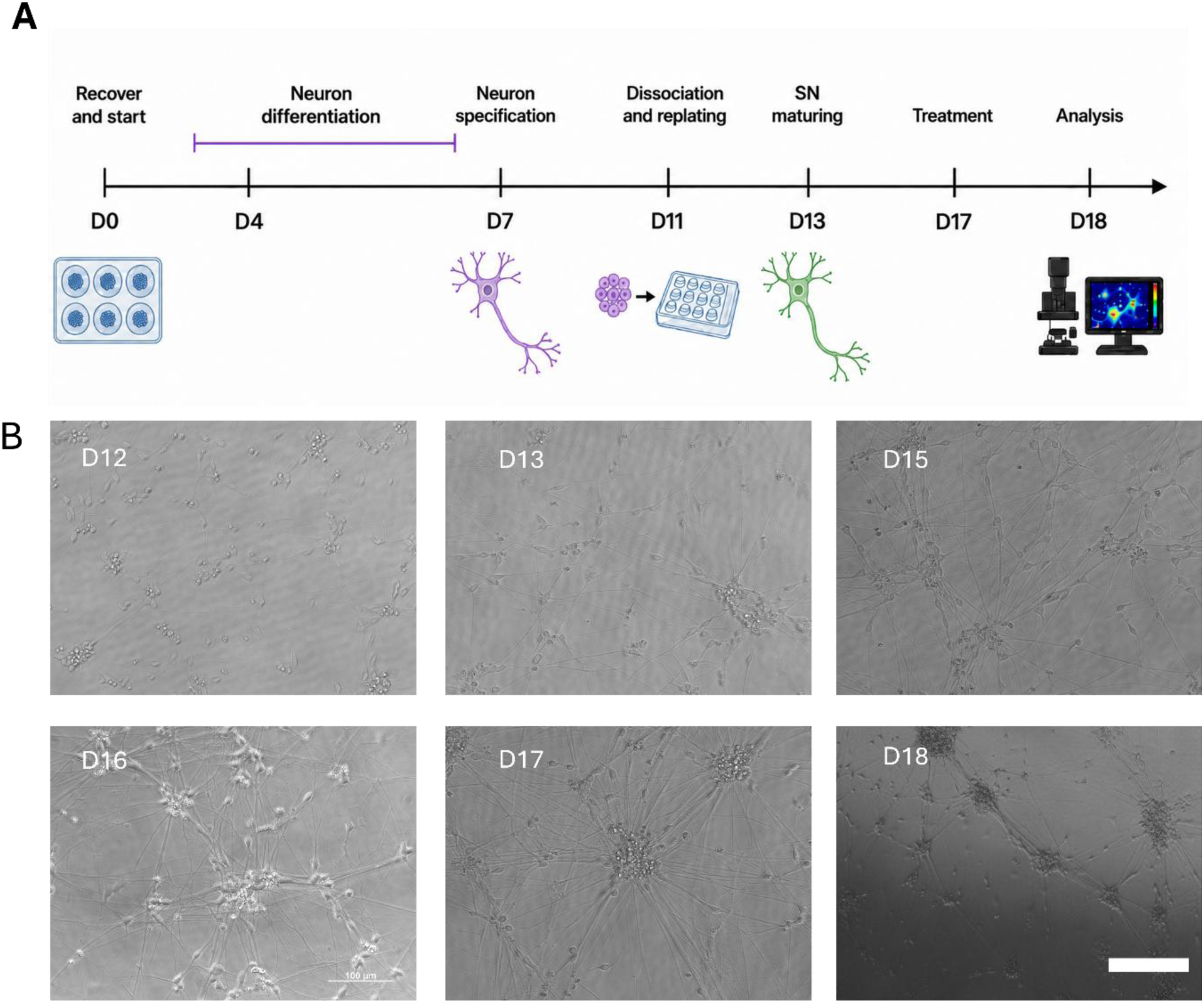
Differentiation of human iPSC-derived sensory neurons and representative bright-field images during neuronal maturation. (A) Schematic overview of the differentiation protocol used to generate human induced pluripotent stem cell-derived sensory neurons (iPSC-SNs), highlighting the major stages of neuronal differentiation from recovery (Day 0) to functional analysis (Day 18) (upper panel). (B) Representative bright-field images illustrating neuronal morphology during maturation following dissociation and replating (Days 12–18) are shown in the lower panels. Progressive neurite extension and the formation of interconnected neuronal networks were observed throughout the maturation period. Scale bar = 200 μm.

**Figure S7.**
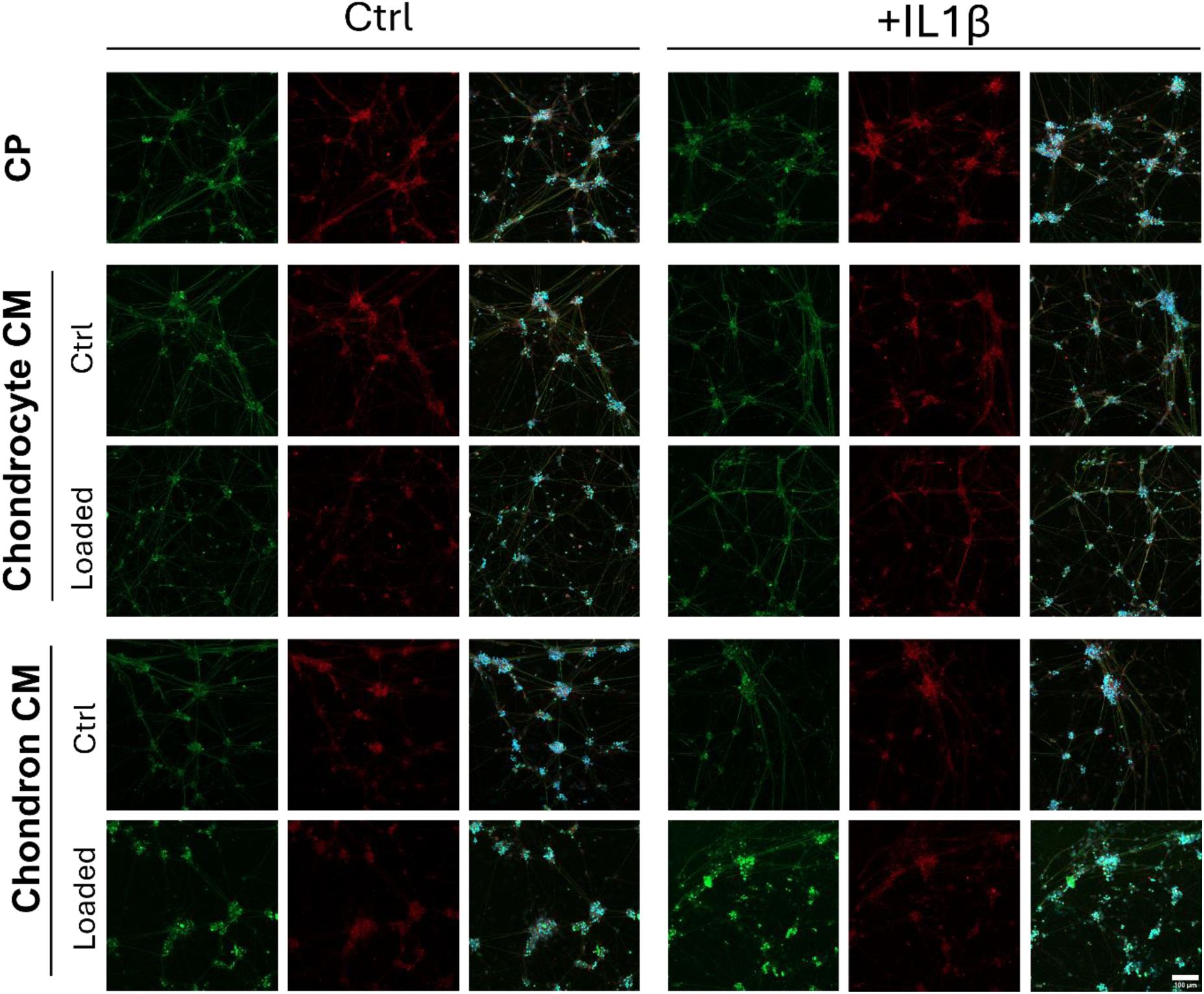
Immunofluorescence characterization of sensory neurons of all conditions. Representative immunofluorescence images of iPSC-derived sensory neurons cultured under all experimental conditions following treatment with conditioned media, and media ctrls (CP). Neuronal networks were stained for βIII-tubulin (green) and calcitonin gene-related peptide (CGRP) (red), with cell nuclei counterstained with DAPI (blue). Merged images demonstrate the maintenance of neuronal morphology and expression of sensory neuronal markers across all treatment groups. Scale bar = 100 μm.

**Figure S8.**
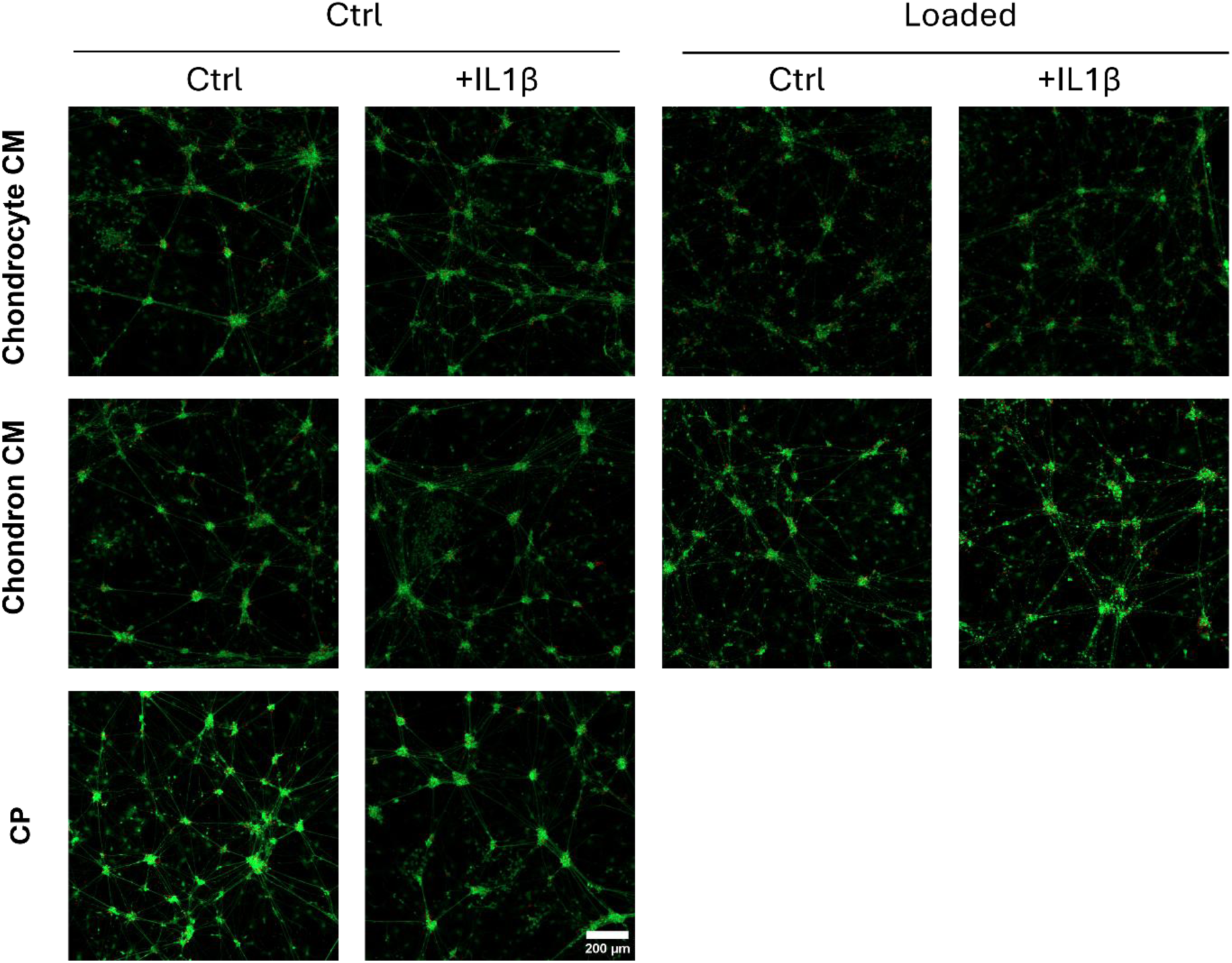
Representative live/dead staining of human iPSC-derived sensory neurons under all experimental conditions. Representative images of human iPSC-derived sensory neurons following treatment with conditioned media from all experimental groups and media ctrls (CP). Cell viability was assessed using a Calcein AM/Ethidium Homodimer-1 (EthD-1) Live/Dead assay, in which viable cells are stained green (Calcein AM) and dead cells with compromised membrane integrity are stained red (EthD-1). Scale bar = 200 μm.

**Figure S9.**
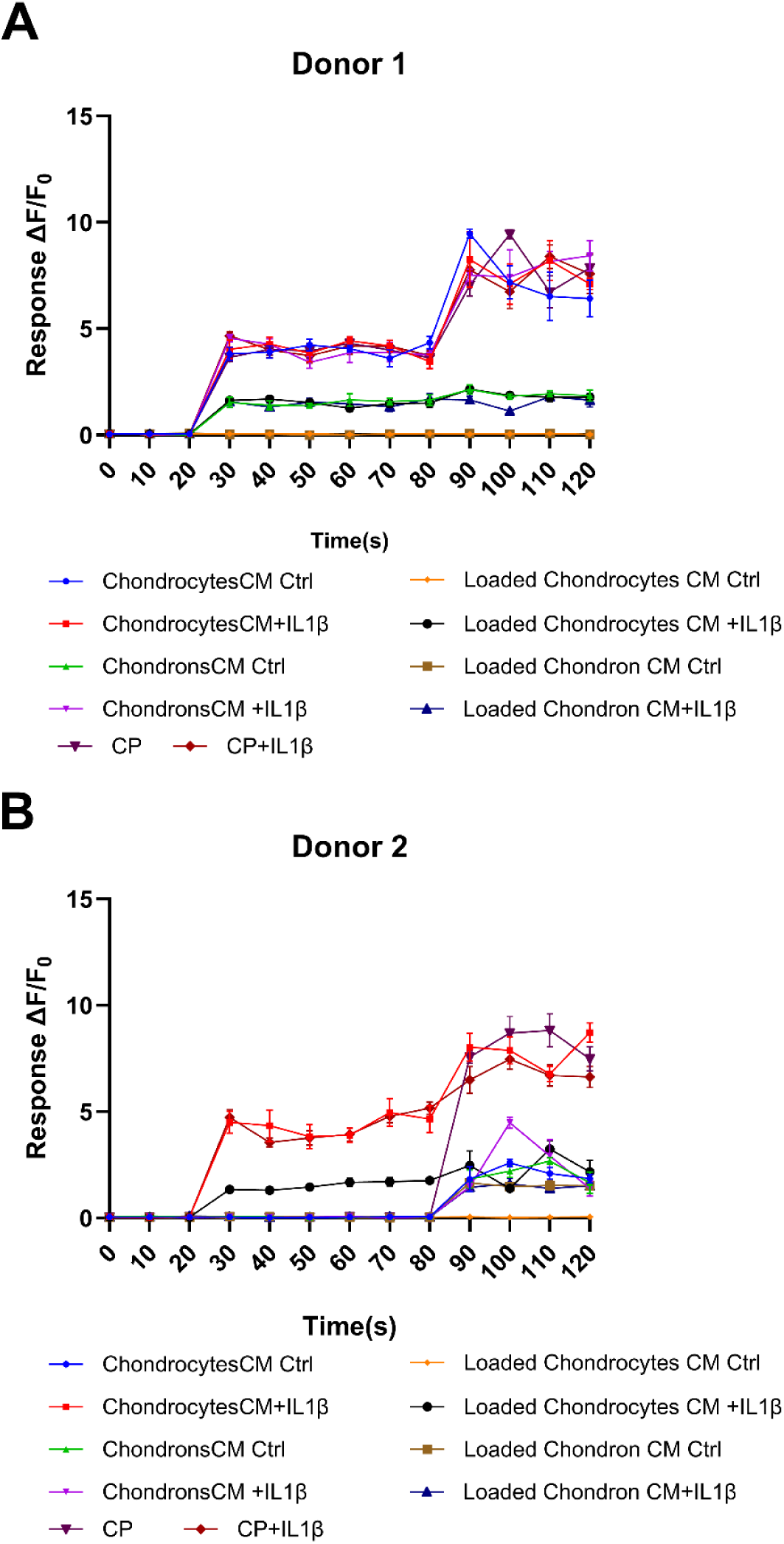
Calcium influx traces of sensory neurons from all conditions. Representative intracellular calcium response traces recorded from human iPSC-derived sensory neurons following exposure to conditioned media from all experimental groups. Responses are shown as changes in Fluo-4 fluorescence intensity (ΔF/F₀) over time for Donor 1 (A) and Donor 2 (B). At 30 s, neurons were stimulated with 100 nM capsaicin to assess TRPV1-mediated calcium responses. At 90 s, neurons were stimulated with KCL. These representative traces correspond to the quantitative analysis presented in Figure 7.

---

**Table S1.**
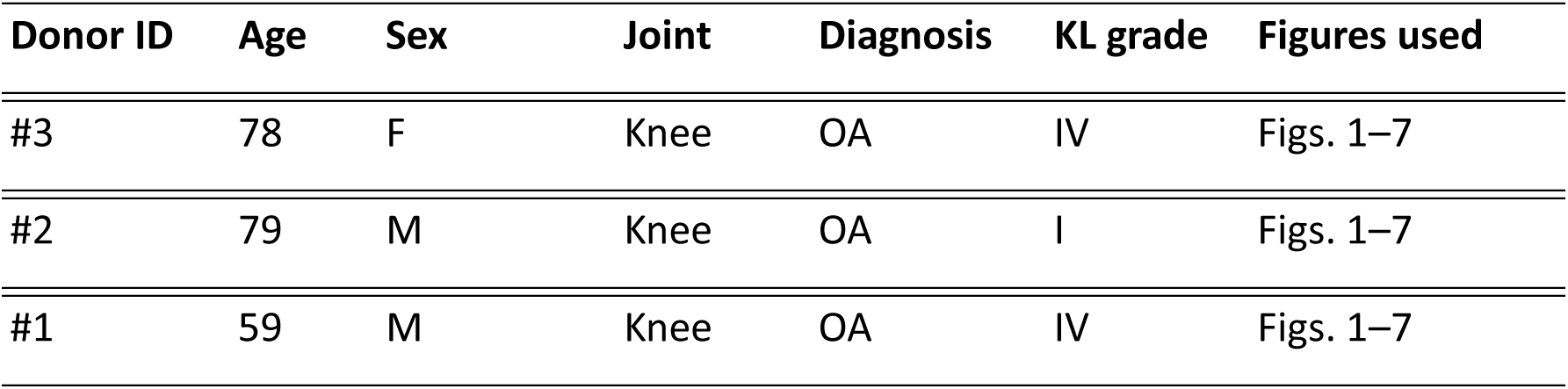
Donor information of chondrocytes sources.

**Table S2.** Antibodies and fluorescent dyes used for immunofluorescence staining.

| Target / antibody | Host | Clone / clonality | Supplier | Catalogue No. | Dilution | Application |
| --- | --- | --- | --- | --- | --- | --- |
| Collagen VI | Rabbit | EPR17072, monoclonal | Abcam, Cambridge, UK | ab182744 | 1:200 | Primary antibody |
| Calcitonin gene-related peptide (CGRP) | Mouse | Clone 5, monoclonal | Thermo Fisher Scientific, Waltham, MA, USA | ABS 026-05-02 | 1:200 | Primary antibody |
| βIII-Tubulin | Rabbit | Polyclonal | Thermo Fisher Scientific, Waltham, MA, USA | PA5-85639 | 1:200 | Primary antibody |
| Goat anti-Rabbit IgG (H+L), Cross-Adsorbed, Alexa Fluor™ 488 | Goat | Polyclonal | Thermo Fisher Scientific, Waltham, MA, USA | **A-11008** | 1:2000 | Secondary antibody |
| Goat anti-Mouse IgG (H+L), Cross-Adsorbed, Alexa Fluor™ 647 | Goat | Polyclonal | Thermo Fisher Scientific, Waltham, MA, USA | **A-21235** | 1:2000 | Secondary antibody |

**Table S3.**
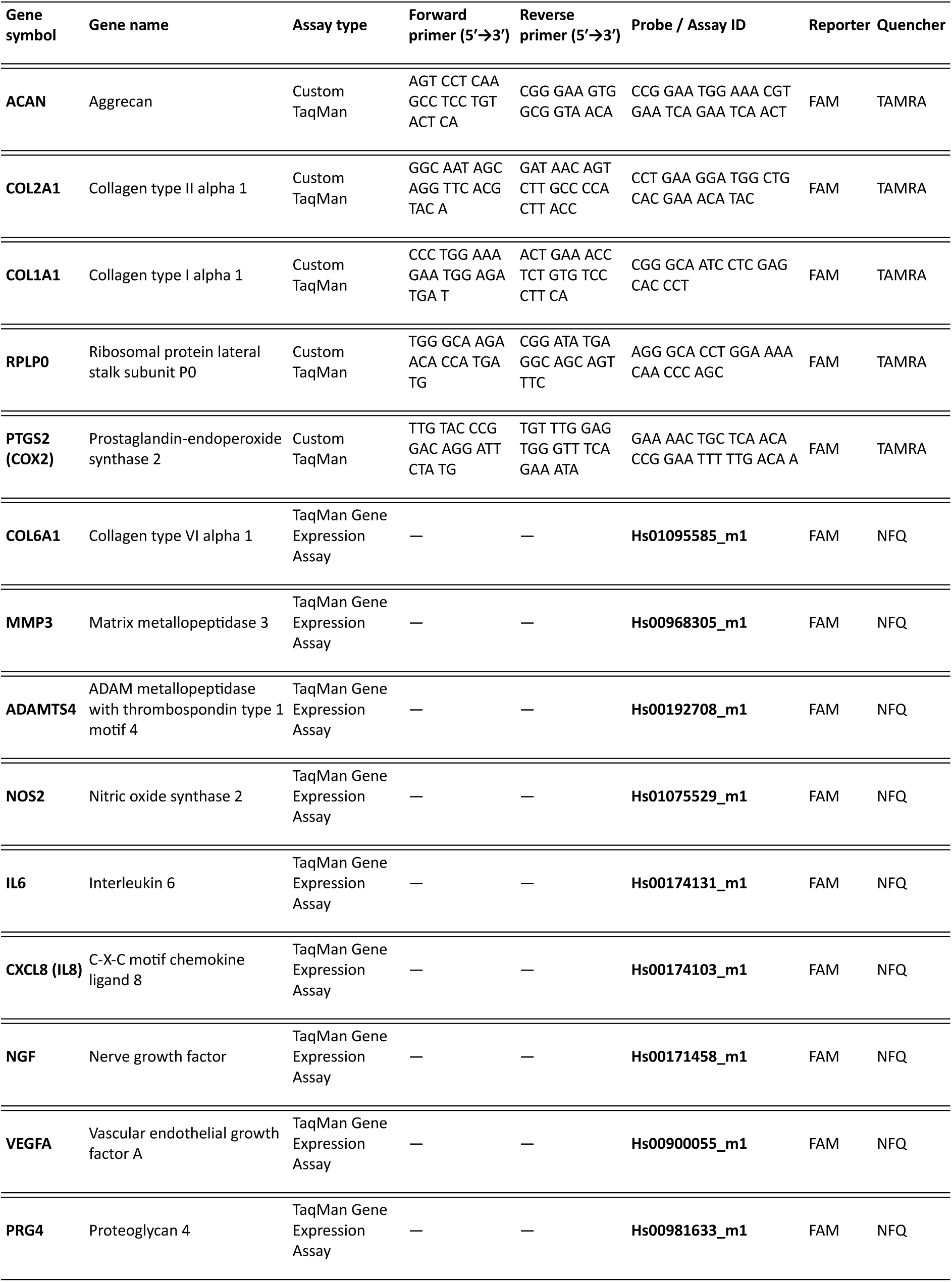
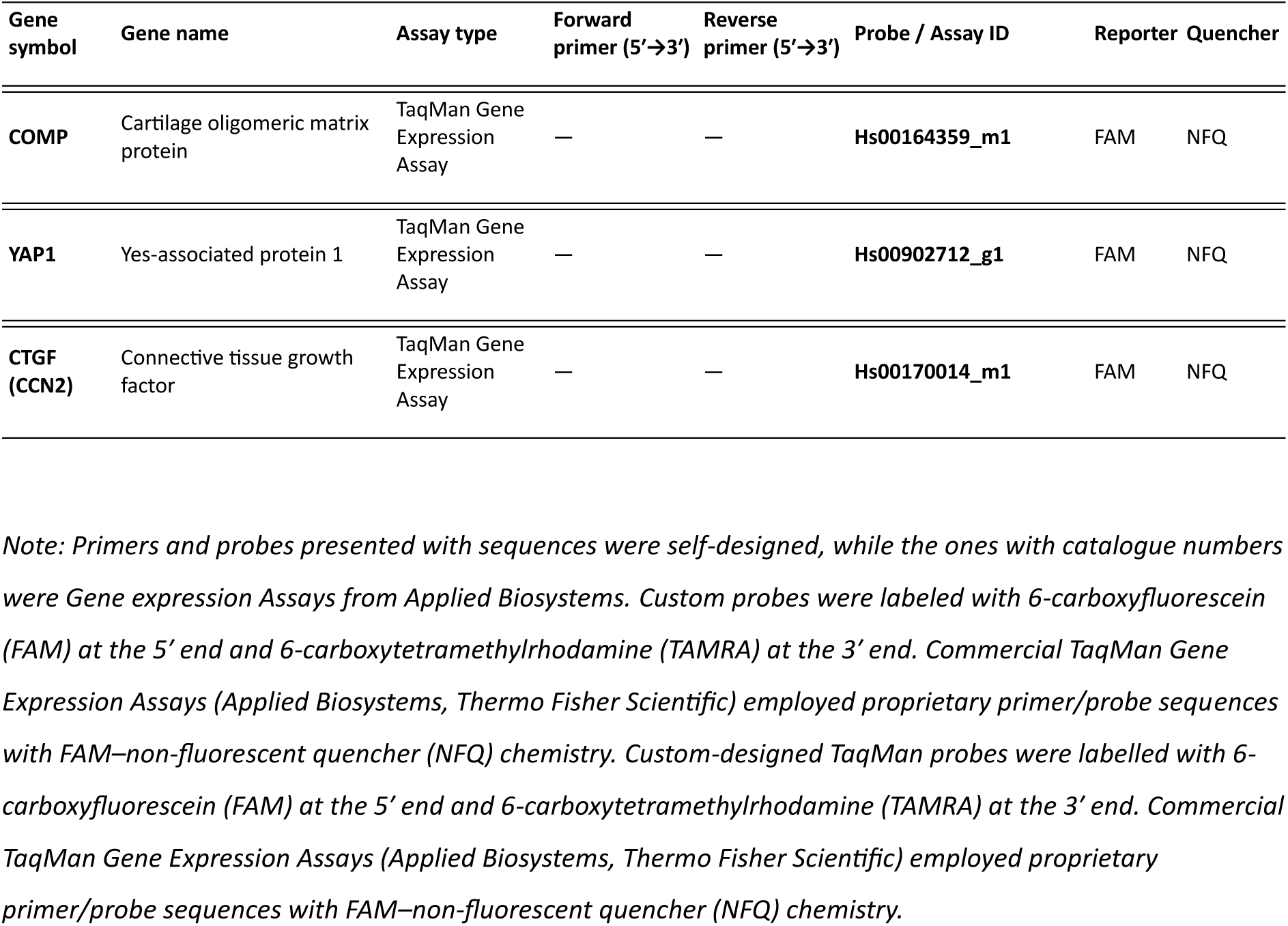
Sequence and source of oligonucleotide primers and probes utilized in RT-qPCR.

**Table S4.** Stage-specific medium composition used for differentiation of human induced pluripotent stem cells into sensory neurons.

| Component | D0–D4 | D5–D6 | D7–D8 | D9–D10 | D11–D12 | D13–D17 |
| --- | --- | --- | --- | --- | --- | --- |
| DMEM/F12 : Neurobasal | 50:50 (v/v) | 50:50 (v/v) | 50:50 (v/v) | 50:50 (v/v) | 50:50 (v/v) | 50:50 (v/v) |
| N2 supplement | 1% v/v | 1% v/v | 1% v/v | 1% v/v | 1% v/v | 1% v/v |
| B27 supplement | 2% v/v | 2% v/v | 2% v/v | 2% v/v | 2% v/v | 2% v/v |
| β-Mercaptoethanol | 0.01 mM | 0.01 mM | 0.01 mM | 0.01 mM | – | – |
| L-Ascorbic acid | 50 µg/mL | 50 µg/mL | 50 µg/mL | 50 µg/mL | 50 µg/mL | 50 µg/mL |
| Y-27632 | 5 µM | 5 µM | 5 µM | 5 µM | 5 µM | 5 µM |
| Glutagro | 1% v/v | 1% v/v | 1% v/v | 1% v/v | – | – |
| MEM non-essential amino acids | 1% v/v | 1% v/v | 1% v/v | 1% v/v | – | – |
| Trace Element A | 0.1% v/v | 0.1% v/v | 0.1% v/v | 0.1% v/v | – | – |
| Trace Element B | 0.1% v/v | 0.1% v/v | 0.1% v/v | 0.1% v/v | – | – |
| Trace Element C | 0.1% v/v | 0.1% v/v | 0.1% v/v | 0.1% v/v | – | – |
| SB431542 | 40 µM | – | – | – | – | – |
| LDN193189 | 0.2 µM | – | – | – | – | – |
| CHIR99021 | 3 µM | 3 µM | – | – | – | – |
| DAPT | – | – | – | 10 µM | 10 µM | – |
| BDNF | – | – | – | – | 20 ng/mL | 20 ng/mL |
| GDNF | – | – | – | – | 10 ng/mL | 10 ng/mL |
| NGF | – | – | – | – | – | 10 ng/mL |
| Gentamycin | 0.1% v/v | 0.1% v/v | 0.1% v/v | 0.1% v/v | 0.1% v/v | 0.1% v/v |

